# Transcriptomic responses of gecarcinid land crabs to acute and prolonged desiccation stress

**DOI:** 10.1101/2024.07.03.601969

**Authors:** Victoria M. Watson-Zink, Richard K. Grosberg, Joelle C.Y. Lai, Rachael A. Bay

## Abstract

Decapod crabs have repeatedly and convergently colonized land. Because of their aquatic ancestry, desiccation is their greatest physiological challenge, yet the genetic basis of their responses to desiccation are unknown. For this study, we sought to identify osmoregulatory genes that were differentially expressed in their antennal glands and posterior gills in response to desiccation stress. We dehydrated and tracked gene expression across three confamilial species displaying increasing degrees of terrestrial adaptation: *Tuerkayana celeste*, *T. magna*, and *Gecarcoidea natalis*. We observed acute dramatic upregulation in the posterior gills of *T. celeste* and *G. natalis* and a more muted response in *T. magna*; however some genes with known osmoregulatory functions were downregulated throughout the trial. We also found that some modules of orthologous genes with correlated expression were associated with greater degrees of terrestriality whereas others reflected shared ancestry, suggesting that different parts of the transcriptome are under varying degrees of terrestrial selective pressure. Finally, while differentially expressed genes were likely to be conserved across the three species, genes from expanded gene families and species-specific genes may also play a role in how land crabs adapt to the unique selective challenges that accompany a terrestrial life.

## 1. Introduction

Most of Earth’s macroscopic biodiversity currently exists on land, but all life began in the oceans (Grosberg et al. 2012). However, sea-to-land transitions are rare, and virtually all extant terrestrial species can be traced to a few ancestral marine lineages (*e.g*., plants, arthropods, vertebrates, and mollusks) that have successfully colonized land and diversified there (Vermeij and Dudley 2000; Vermeij and Watson-Zink 2022). This rarity was decisive in determining Earth’s biodiversity patterns and suggests that this transition presents major physical and physiological challenges that alter nearly every aspect of an animal’s biology. However, little is known about the genomic mechanisms used to overcome these challenges, or how the regulation and function of specific genes shifted as lineages transitioned onto land.

Previous work on the genetic basis of terrestrial adaptation revealed important commonalities between successful terrestrial colonizers, mainly that across the Tree of Life, changes in cellular energy production (Romero et al. 2016), water retention and conservation (Finn et al. 2014; Martinez-Redondo et al. 2023), and osmoregulation and excretion, aerial vision, and oxygen storage and transport (Li et al. 2018; You et al. 2014) enabled this transition. Mechanistically, these studies also suggest that both parallel expansion of ancient gene families and convergently evolved, lineage-specific gene innovations may have facilitated the transition between marine and terrestrial habitats (Aristide & Fernandez 2023). Many of these studies examined lineages that colonized land long ago, such as gastropods, which began colonizing land 307-299 Ma (Vermeij and Dudley 2000), or vertebrates, which began colonizing land 375 Ma (Shubin 2013), obscuring the genetic signatures of these adaptations due to the cumulative effects of neutral evolutionary processes (e.g., relaxed selection on the specific phenotype under consideration, genetic drift, etc.). Yet other studies examine singleton lineages that display some terrestrial tendencies (e.g. mudskippers (You et al. 2014) and walking catfishes (Li et al. 2018)); however, these species lack the phylogenetic counterparts that make comparative phylogenomic approaches possible.

Decapod land crabs are an exceptional example of an ancestrally marine lineage that has recently and repeatedly transitioned into terrestrial habitats. At least seventeen lineages, to varying degrees, have convergently colonized land from the sea over the past 100 million years (Wolfe et al., 2023), either directly via marine-associated terrestrial habitats or after having first adapted to freshwater environments, and each lineage has independently evolved similar solutions to the demands of terrestrial life (Watson-Zink 2021). Due to the vast differences between air and water as media, land crabs have evolved a stunning suite of physiological, morphological, and behavioral strategies to address the selective challenges they encountered while colonizing terrestrial environments, among these being challenges related to osmotic and ionic regulation (Weihrauch et al. 2004), nitrogenous waste excretion (Bliss et al. 1968), aerial respiration (Morris 2002), desiccation and thermoregulation (Wood et al. 1986), growth and molting (Taylor and Kier, 2006), reproduction and development (Anger et al. 2015), visual and olfactory signal transmission and reception (Stensmyr et al. 2005; Krieger et al. 2015), locomotion (Dunham and Gilchrist, 1988; Herreid and Full, 1988; Taylor 2018), and feeding (Greenaway and Linton 1995).

Because terrestriality is a multivariate trait, and crabs exist at varying points along a gradient of terrestrial adaptation, many attempts have been made over the past four decades to describe and categorize terrestriality in this clade. Most recently, Watson-Zink (2021) described four distinct transition pathways (TPs) onto land that crabs have taken over the past 66 million years that account for deep phylogenetic differences between Brachyura and Anomura while also incorporating ecological constraints that each lineage would experience if transitioning to terrestrial habitats via marine-associated terrestrial environments (*e.g*., lower intertidal zone, mudflats, sandflats, mangrove forests) or via freshwater environments and estuaries. The framework ultimately proposes six grades of terrestriality across the four TPs that describe observable trait-by-habitat associations, with Grade I containing crabs that occupy habitats with close ties to full seawater and possess adaptive traits that permit survival in those specific habitats, and Grade VI representing crabs that inhabit arid zones and possess traits that enable their survival in those drier habitats. The high degree of functional and ecological trait convergence displayed by distantly related crab species within each grade sets the stage for investigating, step-by-step, the kinds of genomic and transcriptomic changes that have made such a functionally challenging transition possible for land crabs and other terrestrial animals.

Desiccation is arguably the most pervasive stressor land crabs face in terrestrial habitats. Land crabs as a group are restricted to the tropics and subtropics, and because of the high ambient temperatures and humidities of these regions, the crabs exist at or near their upper limit of heat tolerance (Bliss 1968). In these environments, water evaporates readily and rapidly from any moist tissue or surface, so land crabs have evolved a suite of morphological, behavioral, and physiological adaptations to resist drying out (Watson-Zink 2021). Many land crabs have reduced the size, surface area, and quantity of their gills, which are now primarily used for osmoregulation and excretion (particularly the posterior gills), and have also convergently evolved an accessory respiratory organ (i.e., the branchiostegal lung) to breathe (Burggren and McMahon 1988; Farrelly and Greenaway 1992). Land crabs also drink and reprocess their urine to conserve water and salts (Wolcott and Wolcott 1990), although species differ in terms of which organ they use to reprocess their nitrogenous waste (either the posterior gills or the antennal gland) and how this waste is released (either as a gas released across the gills or as a fluid that must be excreted in water) (Dela-Cruz and Morris 1997; Weihrauch et al. 2004).

To date, there have been no studies published examining the role differential gene expression plays in how land crabs adapt and overcome terrestrial stressors. The dearth of publicly available genomic and transcriptomic resources for this clade make commonly used approaches for linking genes to their functions difficult or impractical. Furthermore, because many of these species are unculturable through successive generations in the lab, it is also not possible to use specialized gene validation techniques to ascertain which genotypes are linked to specific adaptive phenotypes. But comparing gene expression patterns in response to a specific stressor across a set of closely related species may be a means of circumventing these issues while still unearthing genes that are associated with particular adaptive responses.

In this study, we therefore first sought to identify the genes associated with desiccation responses in three closely related land crab species in the family Gecarcinidae (which contains the classically described “land crabs”), particularly in terms of how desiccation stress affects osmoregulation and nitrogenous waste excretion in the posterior gills and the antennal gland. We hypothesized that genes that are canonically known to play critical roles in cellular water movement, osmoregulation, nitrogenous waste excretion, and cellular energy production (e.g., sodium-potassium ATPase, carbonic anhydrase, Rh-like protein, sodium bicarbonate cotransporter, aquaporins, ATP synthase, NADH dehydrogenase, etc.) will be differentially expressed in response to desiccation stress in the selected tissues.

To further explore the role(s) specific genes play in terrestrially adaptive gene expression responses, we also investigated whether modules of orthologous genes with correlated expression across the transcriptome were more strongly associated with a species’ relative degree, or “grade”, of terrestrial adaptation (*sensu* Watson-Zink 2021) or with its phylogenetic distance to either of the other two species in this study. While the null hypothesis would propose greater similarities between close relatives due to shared phylogenetic history (*i.e*., evolutionary conservatism and phylogenetic constraint), past studies have found that distantly related species can evolve similar adaptive solutions to shared selective challenges. Examples of adaptive phenotypic convergence among distantly related species occur across the Tree of Life, such as in camera lenses in the eyes of cephalopods and vertebrates (Ogura et al. 2004), coat pigmentation in beach mice (Steiner et al. 2009), gill raker numbers in marine-to-freshwater sticklebacks (Glazer et al. 2014), transcriptomically similar symbiotic bioluminescent organs in squid (Pankey et al. 2014), and cardiac glycoside insensitivity in milkweed-consuming insect specialists (Groen and Whiteman 2021). Therefore, we asked whether crabs displaying greater degrees of terrestrial adaptation will display more similarities in gene expression patterns across gene modules that are critical to their survival in the terrestrial realm, while the rest of their transcriptome reflects their phylogenetic history.

Finally, because gene repertoire evolution (i.e., gains, duplications, and losses) may play a critical role in adaptive evolution (Aristide and Fernandez 2023; Chen et al. 2013), we examined the role of gene repertoire evolution in the desiccation responses of land crabs to determine whether terrestrially adaptive genes would either be novel, species-specific innovations, or genes that descended from ancient gene families that have been duplicated with expanded functions.

To address these three objectives, we measured tissue-specific differential gene expression in two congeneric land crab sister species from different terrestrial grades, *Tuerkayana celeste* (a freshwater-dependent, air breathing terrestrial crab; Grade III) and *T. magna* (an air-breathing terrestrial crab that digs dry burrows in coastal forests; Grade IV), and a confamilial species, *Gecarcoidea natalis* (an air-breathing terrestrial crab that digs dry burrows in coastal forests that perishes after extended periods of partial immersion in water; Grade V) after exposing the crabs to a time series of increasing desiccation stress (Fig. 1A). Congeneric phylogenetic contrasts between land crabs with differing degrees of terrestriality (i.e., between *T. celeste* and *T. magna*, which diverged ∼700,000 years ago (Ng and Shih 2014)) can reveal how responses to a shared stressor are either constrained by shared ancestry or allowed to vary based on their specific terrestrial niche. On the other hand, confamilial phylogenetic contrasts between more distantly related land crabs with similar degrees of terrestriality (i.e., between *T. magna* and *G. natalis*) may inform our understanding of how similarities in terrestrial niches may drive similarities in gene expression responses between species. By identifying the putative functions and evolutionary histories of genes that are involved in the gene expression responses of land crabs to desiccation stress, we can shed light on how land crabs from different terrestrial grades adapt to the unique selective challenges posed by a life on land and explore the potential role of gene expansion and gene novelty in the adaptive responses of land crabs to terrestrial environments.

**Fig. 1:**
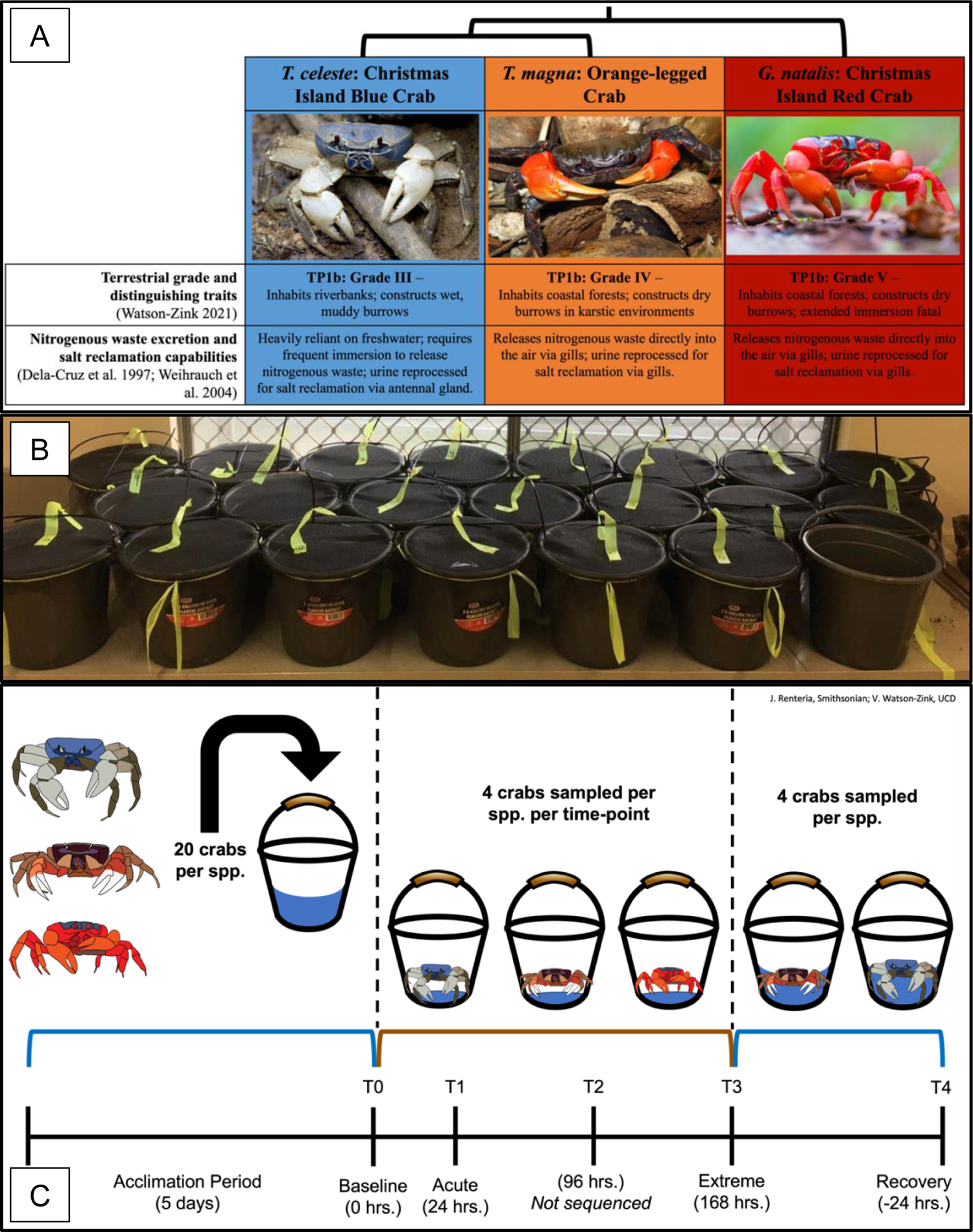
Experimental design and sampling schedule for *T. celeste*, *T. magna*, and *G. natalis*. (A) Cladogram depicting relatedness between three experimental species, their respective terrestrial grades (Watson-Zink 2021), and their distinguishing physiological traits in relation to nitrogenous waste excretion and salt reclamation; (B) Crabs were held in covered 10L buckets throughout the acclimation, desiccation, and recovery periods; (C) Antennal gland and posterior gill 7 tissues were sampled from each crab at four timepoints to measure gene expression responses in each tissue type through time and between species. **Image credits**: “Tuerkayana celeste” by budak is licensed under CC BY-NC SA 4.0. “Indian Ocean Red Claw Land Crab” by Dion Maple is licensed under CC BY-NC 4.0. “Christmas Island Red Crab” by ChrisBrayPhotography is licensed under CC BY-SA 4.0. **Crab pictograph credits**: J. Renteria (Smithsonian) and V. Watson-Zink (Stanford University).

## 2. Materials and Methods

### 2.1 Sample collection and experimental methods

We collected 21 intermolt adult males of *T. celeste* from the stream leading to Anderson’s Dale in Christmas Island National Park, Australia (10°28.7127’ S, 105°33.5138’E) in August 2017, and 21 intermolt adult male *G. natalis* from the dry coastal forest near The Blowholes in Christmas Island National Park (10°51.4458’S, 105°62.7927’E) in September 2017. In December 2017, we obtained 21 intermolt adult males of *T. magna* from the pet trade in Singapore (originally collected in Java, Indonesia). We used 20 individuals from each species for individual expression analyses and one crab per species for *de novo* transcriptome assembly. We collected all Christmas Island specimens by hand and transported them to the laboratory in individual 10L plastic buckets; we also held all *T. magna* specimens in 10L buckets after their arrival from the pet traders (Fig. 1B). We kept all animals at local ambient air temperature (which ranged from 28°C - 41°C) and humidity (which ranged from 60% - 82%) throughout the duration of the acclimation and desiccation treatments.

After selecting only crabs of similar physical condition (i.e., no missing limbs or obvious external physical deformities), we measured each crab’s carapace width (CW) and carapace length (CL) immediately after collection, and weighed each crab twice (once after blotting off excess water before acclimation and again immediately before tissue sampling) to record changes in wet weight throughout the experimental period. On average, *T. celeste* males measured CW: 77.07mm (SD: 7.46mm), CL: 63.55mm (SD: 6.34mm), with initial mean weights of 212g (SD: 57.45g) and ending mean weights of 209g (SD: 56.58g). *G. natalis* males measured CW: 77.36mm (SD: 5.11mm), CL: 57.72mm (SD: 4.21mm), and had initial mean weights of 163.14g (SD: 28.33g) and ending mean weights of 154g (SD: 28.97g). *T. magna* males measured CW: 60.03mm (SD: 4.96mm), CL: 50.13mm (SD: 3.49mm), and had initial mean weights of 104g (SD: 17.93g) and ending mean weights of 102.5g (SD: 16.70g). Because there was very little variation among individual crabs in CW and CL, we assumed that all sampled crabs within each species were approximately at the same developmental stage and could be considered biological replicates. Although all crabs were intermolt, variation in wet weights for similar carapace dimensions could reflect different degrees of intermolt biomass accumulation or hydration status in the wild.

We acclimated all crabs for five days in 10L buckets filled with 1.5L Christmas Island freshwater (CIFW), collected daily from a small stream at Ross Hill Gardens, Christmas Island, Australia (10°29.0923’S, 105°40.8031’E). The water level in the buckets was sufficiently high for a crab to partially submerge its carapace, and therefore all of the posterior gills in its branchial chamber, underwater while sitting; while standing, the water level stood approximately halfway between the crab’s carpi and its propodi (i.e., midway between the second and third segments on the crab’s walking legs). For laboratory studies in Singapore where natural CIFW was not available, we followed the methods of Wood et al. (1986) for imitating freshwater found naturally at the base of land crab burrows on Christmas Island. We refreshed the water in each bucket every 12h. After the 5d acclimation period, we reduced the volume of CIFW in each bucket to 50mL, which was sufficiently high for drinking but not for gill submersion, and maintained the crabs in this desiccation condition for 7 additional days. At the conclusion of the desiccation stress trial, we returned the crabs to acclimation conditions (i.e., 1.5L CIFW in each bucket) to measure their recovery gene expression responses.

Throughout the experiment, we sacrificed four crabs per species at each of five time-points: (1) immediately after the acclimation period ended (0h; “Baseline”); (2) 24h post-acclimation (“Acute”); (3) 96h post-acclimation; (4) 168h post-acclimation (“Extreme”); and (5) 24h after returning all crabs to acclimation conditions (“Recovery”). At each of these time points, we dissected each crab individually on ice. Due to unanticipated mortality during the acclimation period for *G. natalis* and restricted catch limits for this protected species within Christmas Island National Park, we did not sample *G. natalis* for the recovery time point (Fig. 1C). From the crabs sampled for individual gene expression analyses, we took tissue samples from all seven gills from the right side of each crab and its antennal gland; from the one crab per species that we sampled for *de novo* transcriptome preparation, we sampled all seven right-sided gills, the antennal gland, pereiopod muscle tissue, the branchiostegal lung, and the hepatopancreas. We stored all collected samples in 0.5mL cryovials containing Trizol reagent (Invitrogen) and immediately froze them in LN_2._ We transported all samples back to the United States on dry ice held in a 20L dewar storage canister.

### 2.2 Sequencing and transcriptome assembly

We homogenized each tissue sample for one minute in 100µL Trizol reagent (Invitrogen) using a Qiagen Tissue Lyser and beads at 50Hz, and then added an additional 500µL of Trizol to each tube before centrifuging at 16,000G for 30 seconds. We removed the supernatant and placed it into a clean centrifuge tube, and used the Zymo Direct-zol RNA Miniprep extraction kits to extract and purify whole RNA from each sample, following the manufacturer’s instructions. We measured sample concentration and purity using the Agilent 2100 Bioanalyzer (Agilent RNA Eukaryote Total RNA Nano Kit), and the Invitrogen Qubit RNA Broad Range Assay Kit. We then sent all samples to the University of California, Davis Genome and Biomedical Sciences Facility for library preparation and Illumina HiSeq 4000 sequencing. Samples destined for transcriptome assembly were paired-end sequenced at the length of 150bp, while samples slated for differential expression analyses were prepared using a RNA 3’ Tag-Seq protocol and sequenced as single-end 90bp reads.

We checked and visualized read quality using the default parameters in FastQC (Andrews 2010) and eliminated low quality reads with Trimmomatic version 0.36 (Bolger et al. 2014) using parameters for paired reads. We removed leading and trailing low-quality base pairs (LEADING:2, TRAILING:2) using a sliding window of 4 base pairs, removing base pairs when quality dropped below 2 (SLIDINGWINDOW:4:2), and kept reads >25bp (MINLEN:25). In the absence of reference genomes for all three species, we used Trinity (Grabherr et al. 2011) to perform *de novo* transcriptome assemblies for each species using default parameters, and matched all transcripts to NCBI’s BLAST database of protein and translated DNA sequences using DIAMOND (Buchfink et al. 2021) for functional annotation of candidate genes. We matched the top hit per transcript based on e-value and overall percent sequence identity. We then assessed the quality of each assembly with BUSCO’s Arthropoda database (Manni et al. 2021).

### 2.3 Differential gene expression

We estimated transcript abundance using Salmon (Patro et al. 2017), and used DESeq2 (Love et al. 2014) to identify differentially expressed transcripts across three pairwise time intervals: (1) Baseline – Acute, (2) Acute – Extreme, and (3) Extreme – Recovery for all within-species and tissue combinations (a total of six comparisons each for *T. celeste* and *T. magna*, and four comparisons for *G. natalis*). Each time-point consisted of four biological replicates per species. After selecting only genes with *≥* 10 raw reads in *≥* 3 individuals in *≥* 1 of the time-points to filter out genes with low expression, we considered any genes with absolute log2FoldChange expression > 1 and an adjusted p-value < 0.05 to be significantly differentially expressed. For significantly differentially expressed genes, we used a Kruskal-Wallis analysis to determine if the magnitude of differential expression changed across the three pairwise time intervals for each tissue type for all species (total number of analyses = 6). When Kruskal-Wallis results were significant, we used Dunn’s Post-hoc Test with a Bonferroni correction to adjust for multiple comparisons to reveal which time intervals for each tissue type were statistically significantly different. To determine the putative functions of significantly differentially expressed genes, we used a custom Python script to match all DEGs from each species to their respective DIAMOND database entries, and then selected the top 10 genes with the greatest magnitude of differential expression for each time interval for closer examination.

### 2.4 Co-expression analysis

To examine shared and divergent evolutionary patterns of gene expression, we conducted a weighted gene correlation network analysis (WGCNA; Langfelder and Horvath 2008). First, we used TransDecoder (Hass et al. 2013) to translate transcripts into protein sequences and identify putative coding regions, and then used OrthoFinder (Emms and Kelly 2019) to sort the filtered transcripts into orthogroups based on orthologous regions.

After matching single-copy (1:1:1) orthologs shared by all three species with their respective normalized transcript abundance data from Salmon, we then used WGCNA to identify clusters of genes that showed correlated expression patterns across all three species, all four time-points, and both tissue types. We assessed the scale-free topology of our data to choose an appropriate soft-thresholding power (soft thresholding power = 8) by which to raise pairwise correlations of expression, and then transformed ortholog co-expression similarity into a signed adjacency matrix based on the soft thresholding power, which clustered highly correlated genes into modules (minimum module size = 30). We then merged modules with eigengene expression values that were highly correlated (dissimilarity threshold = 0.25), and separated eigengene expression values by tissue type for downstream analysis.

Using a custom Python script, we then matched each ortholog with its respective Trinity transcript identifiers, and used each sequence’s corresponding DIAMOND database output to link each ortholog with an NCBI accession ID based on percent sequence similarity and e-value. From each module, we then selected the top ten genes with the highest absolute module membership (|MM|) to characterize each module’s putative function.

### 2.5 Evolutionary history of differentially expressed genes

To determine if differentially expressed genes (DEGs) were more likely to be assigned to shared orthogroups, and therefore conserved across lineages, we used a custom Python script to match transcript names from DESeq2’s list of significantly differentially expressed genes with transcripts assigned to orthogroups by OrthoFinder. Since OrthoFinder also outputs the copy number for each orthogroup, we also determined if assigned genes were more likely to be single-copy genes or part of expanded gene families (i.e., represented by more than one transcript, although we did not differentiate between isoforms for this analysis). We used Fisher’s exact test to determine if associations between DEGs and one of three gene types (i.e., “unassigned”, “single-copy”, and “expanded”) were more likely for all species, tissue type, and time-point combinations (total number of Fisher tests = 32), and then ran a Kruskal-Wallis analysis to determine if there were differential expression differences between gene type categories for all species, tissue type, and time-point combinations (total number of analyses = 16). If prior Kruskal-Wallis results were significant, we then used Dunn’s Post-hoc Test with a Bonferroni correction to adjust for multiple comparisons to determine whether there were statistically significant differences between the differential expression of pairwise gene type groupings for each time interval.

## 3. Results

### 3.1 *de novo* reference transcriptome assembly

We obtained 89 – 150 million raw reads per transcriptome, which were assembled into about 113 – 221 thousand transcripts per transcriptome (Table 1). Mean contig N50 was 1,482 base pairs, and the mean GC content for each transcriptome was 46%. The BUSCO Arthropoda database revealed that the *T. magna* transcriptome was the most complete (88% complete), followed by the *T. celeste* transcriptome (82% complete) and the *G. natalis* transcriptome (79% complete). Since the BUSCO Arthropoda database was compiled using mostly terrestrial arthropods (i.e., insects and arachnids), it is not surprising that many transcripts were not represented in any of the decapod crustacean transcriptomes compiled in this study.

**Table 1:** *de novo* transcriptome summary statistics for the three species represented in this study.

| *T. celeste* |  | *T. magna* |  | *G. natalis* |  |
| --- | --- | --- | --- | --- | --- |
| Total transcripts | 221,111 | Total transcripts | 143,688 | Total transcripts | 113,191 |
| Total bases | 150,915,012 | Total bases | 133,242,897 | Total bases | 89,956,155 |
| GC% | 47% | GC% | 46% | GC% | 46% |
| BUSCO<br>(Arthropoda) | 82% | BUSCO<br>(Arthropoda) | 88% | BUSCO<br>(Arthropoda) | 79% |
| **All transcript contigs** |  | **All transcript contigs** |  | **All transcript contigs** |  |
| N50 transcript<br>length (bp) | 1,114 | N50 transcript<br>length (bp) | 1,769 | N50 transcript<br>length (bp) | 1,562 |
| Smallest contig<br>length | 201 | Smallest contig<br>length | 201 | Smallest contig<br>length | 201 |
| Mean contig<br>length | 682.53 | Mean contig<br>length | 927.31 | Mean contig<br>length | 794.73 |
| Largest contig<br>length | 17,698 | Largest contig<br>length | 19,701 | Largest contig<br>length | 21,899 |

### 3.2 Tissue-specific differential gene expression

#### 3.2.1 Tuerkayana celeste

*T. celeste* responds to sudden desiccation stress primarily by dramatically shifting how genes are expressed in its posterior gills, as well as in its antennal gland to a much smaller degree. The majority of DEGs were expressed during the Baseline-Acute time interval in both the posterior gill (3,503 transcripts) and the antennal gland (181 transcripts), and most of these genes were upregulated (Fig. 2A). The dramatically upregulated gene expression response displayed by the posterior gill during the Baseline – Acute time interval, where we recorded several orders of magnitude more DEGs than at any other time interval for this species, was especially notable (Fig. 3A; Fig. 3B). In the posterior gill, there were statistically significant differences in gene expression across all three time intervals, indicating that throughout the time series trial there were large shifts in overall gene expression patterns in this tissue in response to acute and prolonged desiccation stress and eventual re-immersion (Fig. 2B).

**Fig. 2:**
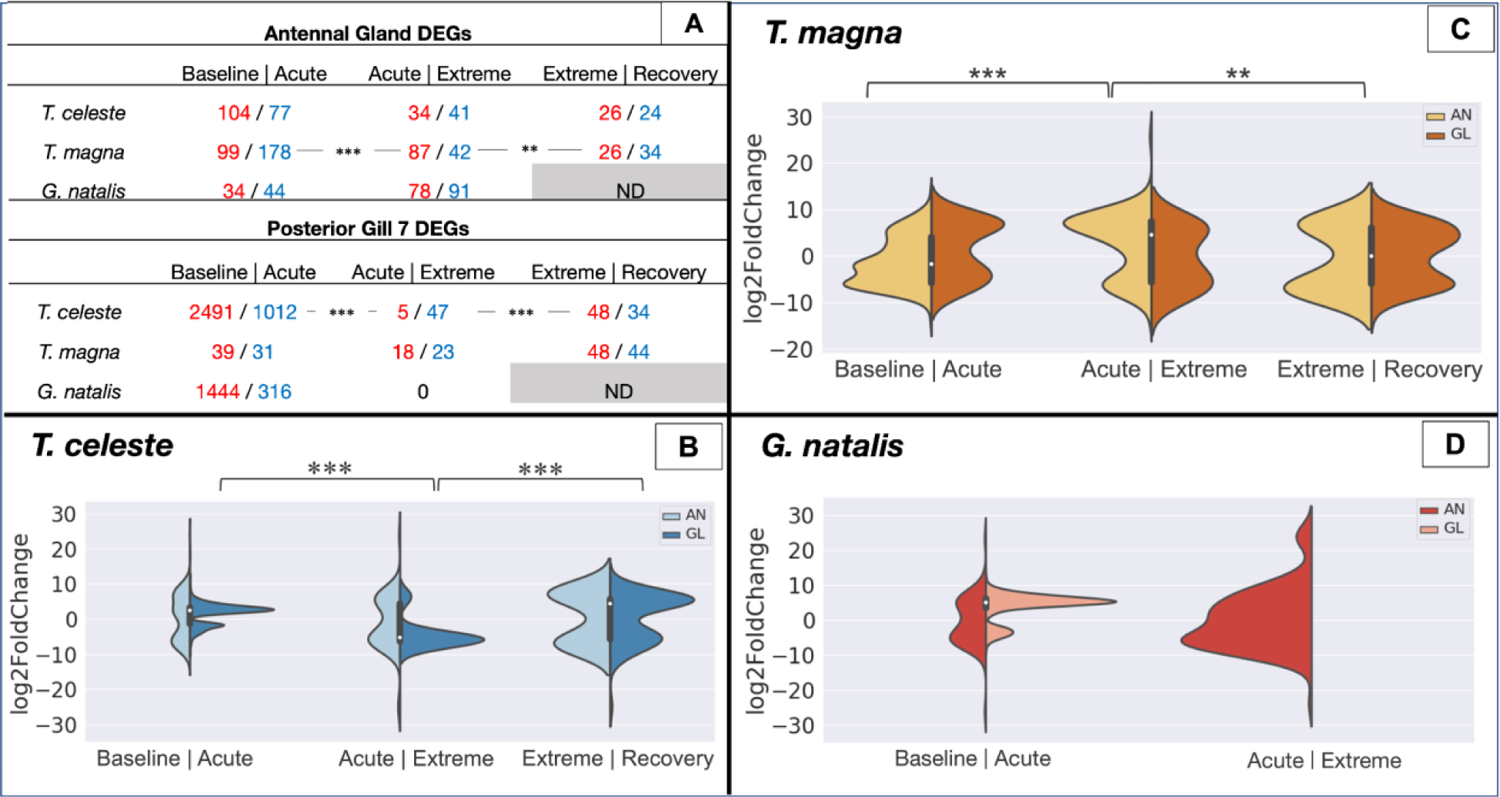
Tissue-specific differentially expressed genes for all three species through time. (A) Table depicting quantities of upregulated (red text) and downregulated (blue text) genes during each time interval for both tissue types and across all three species; (B-D) Violin plots depicting distribution of log2FoldChange expression values and summary statistics during each time interval for both tissue types for *T. celeste* (B), *T. magna* (C), and *G. natalis* (D). Asterisks reflect degree of statistical significance between time interval pairings: p < 0.05 (*); p < 0.01 (**); p < 0.001 (***).

**Fig. 3:**
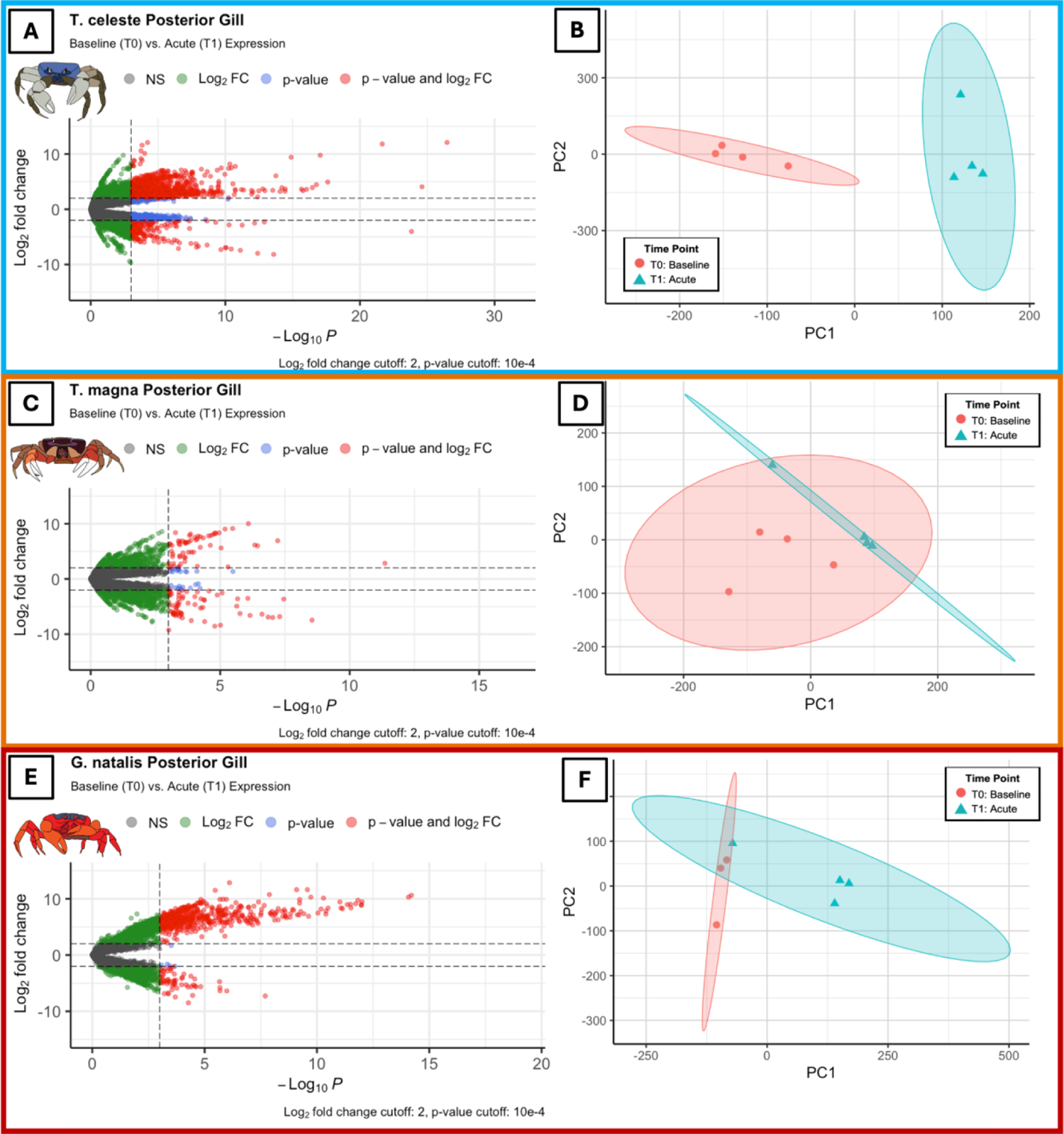
(A, C, E): Volcano plots showing mean of normalized counts by log2FoldChange expression for *T. celeste* (A), *T. magna* (C), and *G. natalis* (E) during the Baseline – Acute time interval. (B, D, F): Principal component analyses between four biological replicates each for *T. celeste* (B), *T. magna* (D), and *G. natalis* (F) during the Baseline – Acute time interval. 95% confidence intervals for each time point are shown in shaded ellipses. **Crab pictograph credits**: J. Renteria (Smithsonian) and V. Watson-Zink (Stanford University).

#### 3.2.2 Tuerkayana magna

The majority of DEGs for *T. magna* occurred in antennal gland tissues during the Baseline – Acute interval, where most of the DEGs were downregulated (277 transcripts total), and in the antennal gland tissues during the Acute - Extreme interval, where most of the DEGs were upregulated (129 transcripts total) (Fig. 2A; Fig. 2C). The signal in the posterior gill during the Extreme – Recovery time interval was elevated but comparatively moderate, where there was an almost equal number of DEGs upregulated as there were downregulated (92 transcripts total) (Fig. 2A). In the antennal gland, there were statistically significant differences in gene expression across all three time intervals, indicating that throughout the time series trial there were significant shifts in overall gene expression patterns in this tissue in response to acute and prolonged desiccation stress and eventual re-immersion (Fig. 2C).

#### 3.2.3 Gecarcoidea natalis

As was the case for *T. celeste*, *G. natalis* also appeared to respond to acute desiccation stress by upregulating genes in its posterior gill. Most DEGs for *G. natalis* were expressed in the posterior gill during the Baseline – Acute time interval (1,760 transcripts); overall, the gene expression signal during this interval was one of significant upregulation (Fig. 2A; Fig. 3E – F). In comparison, during the subsequent time interval (Acute – Extreme), there were no genes that were significantly differentially expressed in the posterior gill (Fig. 2A).

### 3.3 Orthology assignment and co-expression analysis

TransDecoder identified a total of 131,781 coding sequences across all three transcriptomes (52,781 coding sequences in *T. celeste*, 45,084 coding sequences in *T. magna*, and 33,684 coding sequences in *G. natalis*). OrthoFinder assigned 85,186 of these genes (64.8% of total) into 17,661 orthogroups with a mean orthogroup size of 4.8 genes, and a median orthogroup size of 4 genes (although the largest orthogroup consisted of 91 genes). Fifty-eight orthogroups were species-specific (17 were specific to *T. celeste*, 14 were specific to *T. magna*, and 27 were specific to *G. natalis*, while 9,516 orthogroups were present in all three species (Fig. 4). OrthoFinder recovered 2,185 1:1:1 single-copy orthogroups, all of which were used for weighted gene correlation network analyses.

**Fig. 4:**
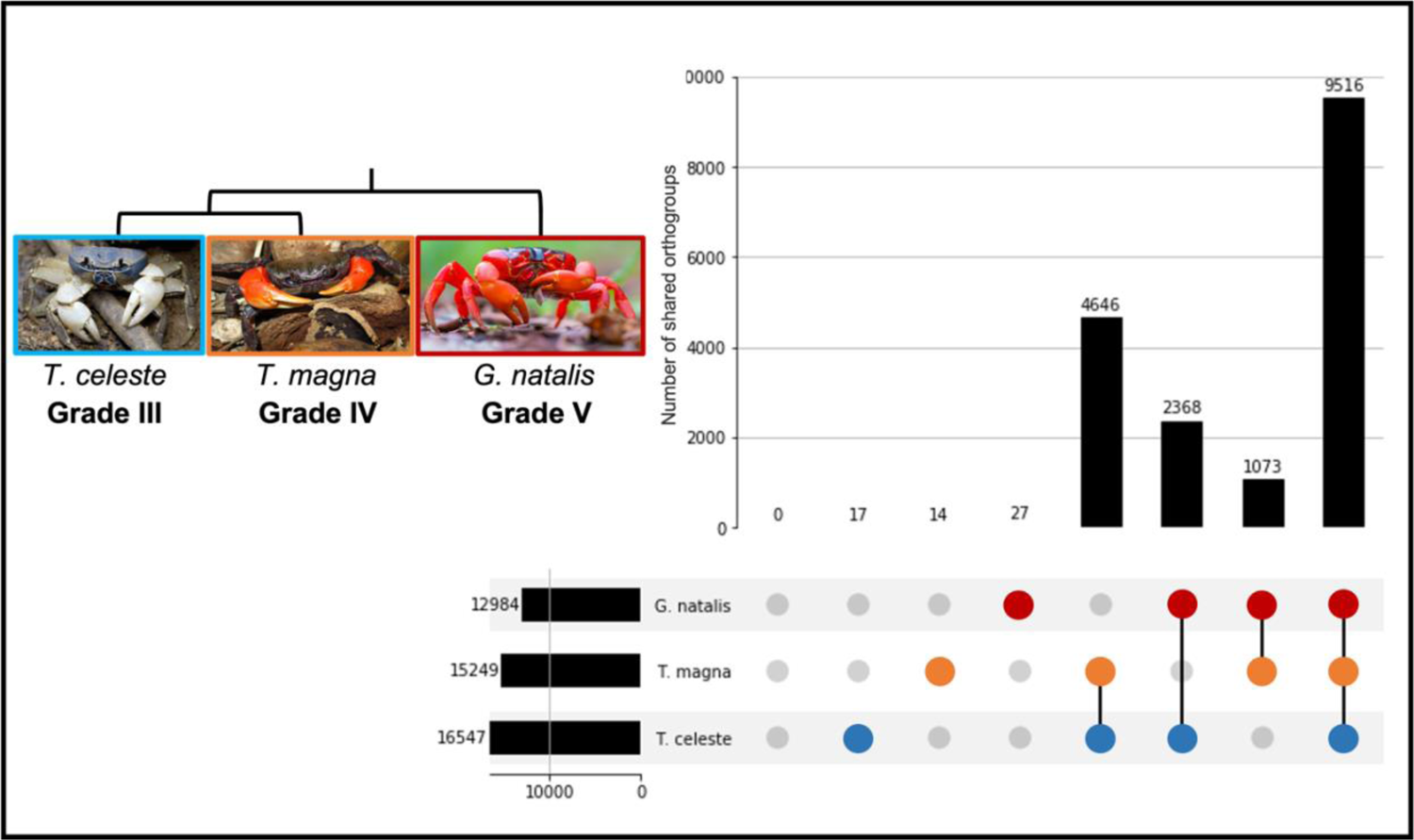
Upset diagram of OrthoFinder results, showing the relative contributions of each species (*T. celeste*, *T. magna*, and *G. natalis*) to the orthogroups, and intersections between species. Each number represents the number of orthogroups unique to that species or grouping of species. A graphical representation of total orthogroups per species is also plotted along the x-axis. Inset: Pictorial cladogram displaying phylogenetic relationships between species and relative terrestrial grades. **Image credits**: “Tuerkayana celeste” by budak is licensed under CC BY-NC SA 4.0. “Indian Ocean Red Claw Land Crab” by Dion Maple is licensed under CC BY-NC 4.0. “Christmas Island Red Crab” by ChrisBrayPhotography is licensed under CC BY-SA 4.0.

After merging modules with highly correlated eigengene values (dissimilarity threshold = 0.25), WGCNA clustered 2,185 1:1:1 orthologs into six modules of highly correlated genes; unclustered genes were assigned to Module 0. Eigengene expression is defined as the first principal component of a given module, and represents the overall gene expression profile of that particular module. We therefore used sample eigengene values to split each module’s eigengene expression into relative contributions by tissue type (i.e., by the posterior gill and the antennal gland) to determine how module expression varied between species through time in each tissue type.

Overall, eigengene expression reflected differences between the three species rather than experimental treatments. Five modules showed statistically significant species associations, while none of the modules were significantly associated with any of the time points (Fig. 5A). Across species, some modules showed greater similarities between congeners in terms of the magnitude and directionality of their gene expression responses (a” phylogenetic effect”; Modules 1 and 3), while other modules displayed a “grade effect”, where expression profiles were more similar across crab species from higher terrestrial grades (Modules 5 and 6). Finally, Module 4 was strongly positively associated with *T. magna* alone; genes in this module were expressed at constitutively lower levels in the remaining two species (Fig. 5B). Module 2 was also weakly negatively associated with *T. magna*, but this pattern was not statistically significant. Finally, all modules showed strongly significant associations across tissue types; these associations most likely resulted from differences in transcript abundance between the tissues.

**Fig. 5:**
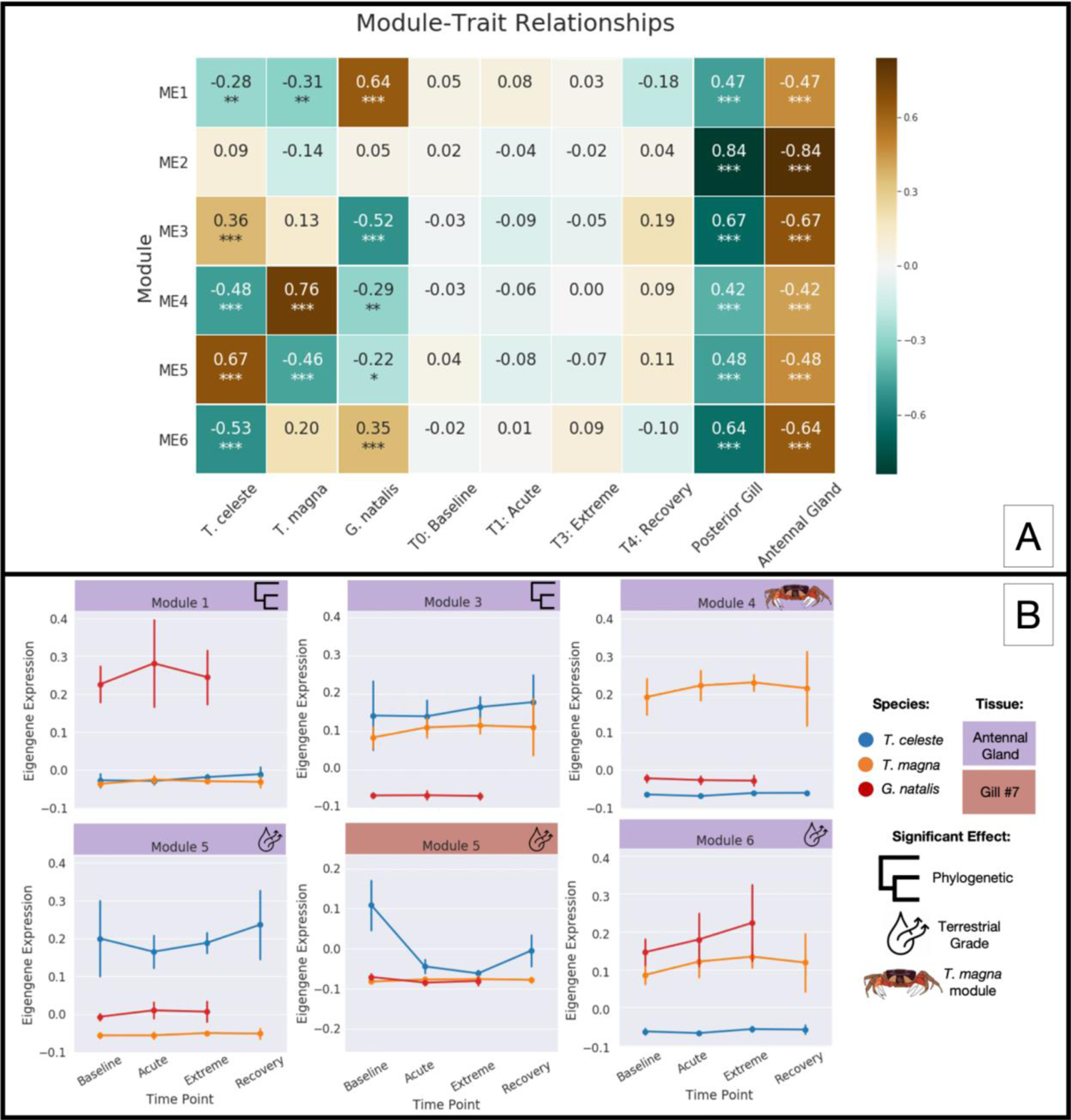
(A) Module-trait relationships of co-expression modules designated by WGCNA (Langfelder and Horvath 2008). Warmer colors represent more positive correlations between particular modules and traits (*i.e*., timepoints, species, tissue types), while cooler colors represent more negative correlations between the aforementioned factors. Asterisks reflect degree of statistical significance between relationships: p < 0.05 (*); p < 0.01 (**); p < 0.001 (***). (B) Tissue-specific eigengene expression values for key co-expression modules for all three species. Icons depict significant effects for each module (*i.e*., phylogenetic effects, grade effects, etc.). Only modules with statistically significant associations are shown.

### 3.4 DEG gene repertoire and evolutionary histories

Differentially expressed genes within species tended to be conserved across all three species (i.e. assigned to an orthogroup shared across species) (Supplemental Materials, Table 1), suggesting that gecarcinid land crabs respond to desiccation stress primarily by using conserved genes. Additionally for *T. celeste*, differentially expressed genes in the posterior gill at the Baseline - Acute and Acute - Extreme time intervals also tended to be single-copy genes, and this was also true for differentially expressed genes in the posterior gill of *G. natalis* during the Baseline - Acute time interval. In nearly every case, except for the Baseline – Acute time interval in the posterior gill of *T. celeste* (H(2) = [321.55], p = 1.5 x 10^-70^) and the posterior gill of *G. natalis* (H(2) = [7.39], p = 2.5 x 10^-2^) (Fig. 6), there were no statistically significant differences in gene expression between gene types (i.e., single-copy conserved genes, genes from expanded gene families, and species-specific “unassigned” genes). For the Baseline – Acute time interval in the posterior gill of *T. celeste*, Dunn’s Post-hoc Test showed that mean differential expression significantly differed between unassigned genes and single-copy genes (p = 3.89 x 10^-44^), and between unassigned genes and genes from expanded gene families (p = 2.84 x10^-34^) (Fig. 6). For the Baseline – Acute time interval in posterior gill of *G. natalis*, Dunn’s Post-hoc Test for multiple comparisons showed that mean differential expression significantly differed only between unassigned genes and single-copy genes (p = 2.1 x 10^-2^). These results suggest that while some component of the desiccation stress response in *T. celeste* and *G. natalis* can be traced back to single-copy conserved genes, a significant portion is driven by genes unique to these species and, for *T. celeste*, also by genes that have undergone gene family expansion. Furthermore, the directionality of expression shifts across these categories varied: for *T. celeste*, unassigned genes during the Baseline – Acute time interval were overwhelmingly upregulated, while single-copy genes tended to be downregulated. Genes from expanded gene families appeared to be modestly bimodal – some were upregulated, while others were downregulated (Fig. 6). For *G. natalis*, however, both unassigned genes and single-copy conserved genes tended to be upregulated during the Baseline – Acute time interval.

**Fig. 6:**
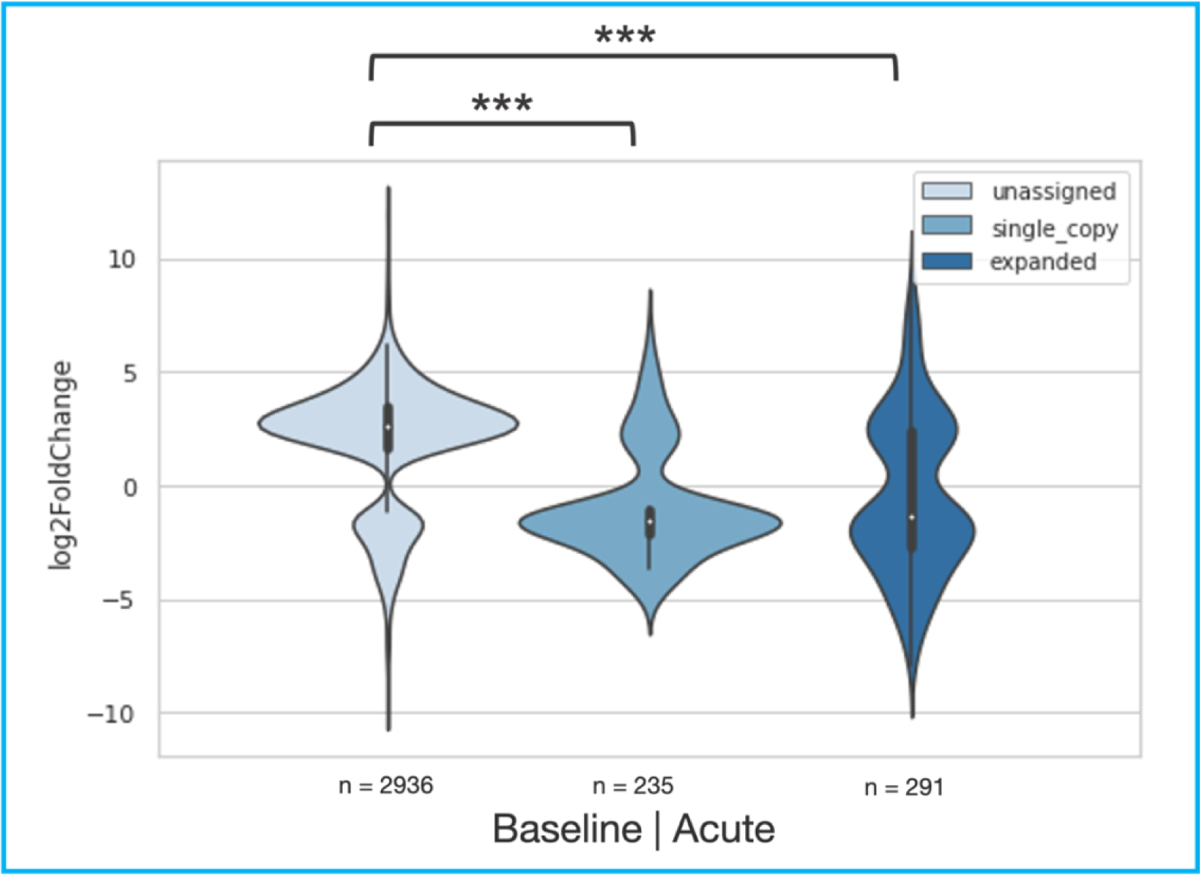
Violin plots representing quantities and log2FoldChange expression values and distributions for different gene types (*i.e*., “unassigned”: genes not clustered in any shared orthogroups by WGCNA; “expanded”: genes having more than one copy in an orthogroup for a single species; “single-copy”: genes that have only one copy in an orthogroup for that species) during the Baseline – Acute time interval in the posterior gill tissue of *T. celeste*. Asterisks reflect degree of statistical significance between log2FoldChange expression values of gene type comparisons: p < 0.05 (*); p < 0.01 (**); p < 0.001 (***).

### 3.5 Candidate gene function of DEGs by species

For *T. celeste*, the majority of the top 10 genes with the greatest magnitude of differential expression between time intervals for each species-by-tissue-by-time interval combination had no corresponding matches in the NCBI BLAST database. Of the genes that successfully matched publicly available BLAST records, none were single-copy conserved genes.

In the posterior gill during the Baseline – Acute time interval, genes related to protein synthesis were upregulated, while genes related to the degradation of ubiquitin fusion proteins were upregulated during the Acute – Extreme time interval. None of the other top 10 genes for any of the time intervals in the posterior gills matched functionally informative records.

In the antennal gland during the Baseline – Acute time interval, protein kinase C, which is responsible for signal transduction pathways in the cell, was upregulated. During the Acute –Extreme time interval in the antennal gland, there was significant downregulation of genes related to steroidogenesis and chloride channel synthesis. Finally, during the Extreme – Recovery time interval in the antennal gland, there was significant upregulation in genes related to *domeless*, which plays a role in border cell migration (Supplemental Materials, Table 2).

**Table 2:**
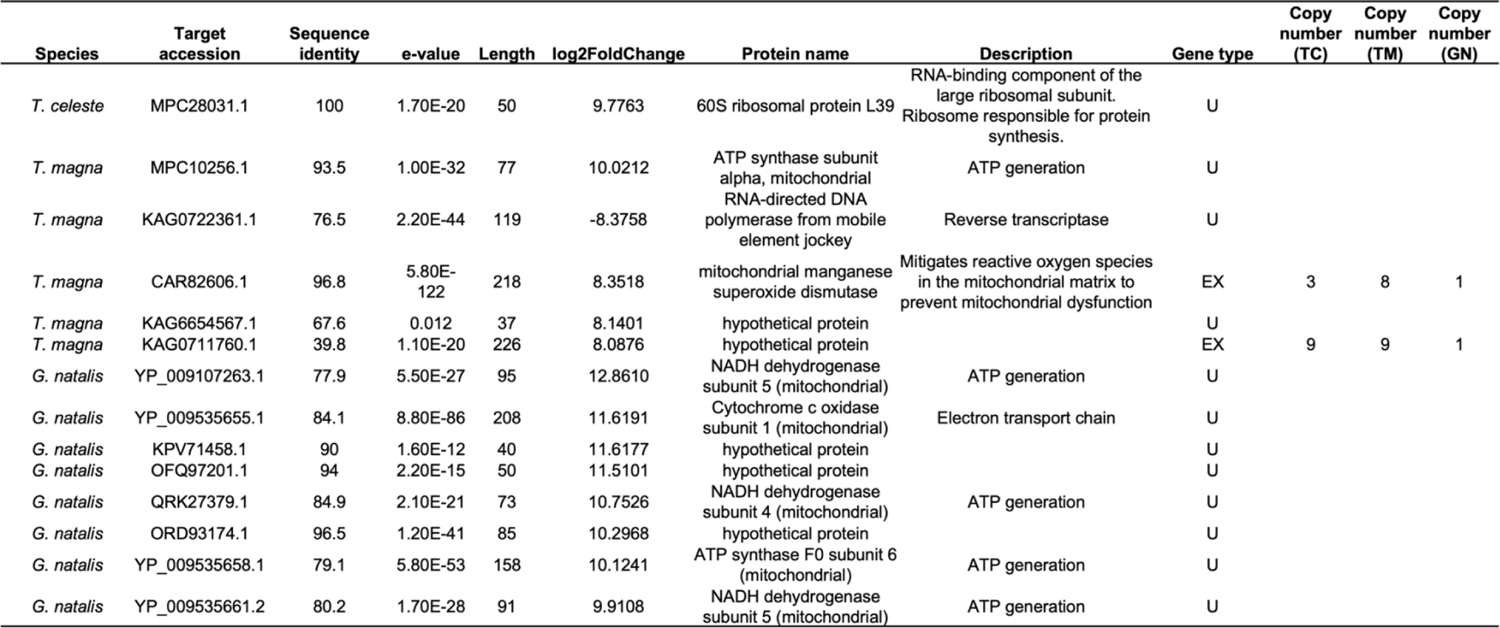
Gene names and descriptions for top 10 DEGs in terms of log2FoldChange expression in the posterior gill for the Baseline-Acute time interval for all three species. Only genes with BLAST hits are shown. Gene types (EX: expanded gene family; U: unassigned to any orthogroup) and copy numbers per species (TC: *T. celeste*; TM: *T. magna*; GN: *G. natalis*) are also shown.

For *T. magna*, many more of the top 10 log2FoldChange genes matched to BLAST entries than for *T. celeste*. As was the case for *T. celeste*, none of these genes were single-copy conserved genes. Overall, there were significant expression changes in genes involved in ATP synthesis, chromatin structure maintenance and remodeling, transmembrane transport, glycolysis, DNA repair, pyrimidine degradation, and in those responsible for protecting the mitochondria from oxidative damage (Supplemental Materials, Table 3). There was also downregulation in two transposon-related proteins (*i.e*. proteins related to *tigger*, which is part of the *pogo* DNA-mediated transposon family, and *jockey*, which encodes an RNA-directed DNA polymerase).

For *G. natalis*, more of the top 10 log2FoldChange genes had corresponding BLAST hits than we observed for *T. celeste*, but there were fewer matches than there were in *T. magna*. As was the case for both the other species, none of these genes were single-copy conserved genes. Overall, genes encoding energy-producing mitochondrial enzymes, including NADH dehydrogenase (subunits 4 and 5), ATP synthase, and Cytochrome C oxidase subunit 1, were significantly upregulated in the posterior gill during the Baseline-Acute time interval. In the antennal gland, genes related to alternative splicing, transcription termination, and the maintenance of pluripotency were significantly upregulated (Supplemental Materials, Table 4).

Finally, genes that are canonically known to play critical roles in cellular water movement, osmoregulation, and nitrogenous waste excretion (e.g., sodium-potassium ATPase, carbonic anhydrase, Rh-like protein, sodium bicarbonate cotransporters, aquaporins, etc.) displayed varying signals across all three species. Of the known osmoregulatory genes that were significantly differentially expressed in this study, the following patterns emerged. In *T. celeste*, throughout the duration of the study, many of these genes were differentially expressed between the Baseline and Acute time points in the posterior gill, where the sodium-potassium ATPase, sodium-bicarbonate cotransporter, and Rh-like protein were all downregulated (log2FoldChange expression of −3.09, −4.02, and −4.00, respectively). Only ankyrin/Trp cation channel was upregulated during this time interval, with a log2FoldChange expression of 2.36 (Supplemental Materials, Figure 1). In *T. magna*, these genes were only differentially expressed either in the posterior gill between the Extreme – Recovery time points (ankyrin-2/Trp cation channel was upregulated: log2FoldChange expression of 8.22), or in the antennal gland between the Baseline – Acute time points (glucosidase was upregulated: log2FoldChange expression of 1.82, as well as the sodium-driven bicarbonate exchanger, which was also upregulated with a log2FoldChange expression of 1.81). In *G. natalis*, these genes tended to be expressed in the gills between the Baseline – Acute time points: ankyrin/Trp cation channel, carbonic anhydrase, and DnaJ subfamily B, which interacts with HSP70, were all upregulated (log2FoldChange expression of 5.90, 3.47, and 5.24, respectively), while aquaporin AQPcic was downregulated with a log2FoldChange expression of −3.58 (Supplemental Materials, Figure 2). In the antennal gland during this same time interval, ankyrin was significantly downregulated with a log2FoldChange expression of −6.34.

### 3.6 Putative functions of single-copy conserved genes

After selecting the top ten 1:1:1 orthologs with the highest absolute module membership for each module and matching these genes to their respective BLAST hits, we found that nearly all of the selected genes encoded specific transcription factors and chaperone proteins that assist in protein folding. There were also several genes that matched enzymes responsible for DNA repair (Supplemental Materials, Table 5).

## 4. Discussion

Terrestrial crabs have independently colonized land to varying degrees at least seventeen times in the past 100 million years (Wolfe et al. 2023). They display a diversity of ecotypes and inhabit a variety of terrestrial landscapes while also showing a high degree of functional and ecological convergence in response to the significant adaptive challenges that accompany a transition to terrestrial life (Watson-Zink 2021).

In this study, we used tissue-specific RNAseq to assess how three gecarcinid land crabs from different terrestrial grades (*Tuerkayana celeste*, Grade III; *T. magna*, Grade IV; and *Gecarcoidea natalis*, Grade V) responded to desiccation, their most pervasive terrestrial stressor, across the posterior gills and the antennal gland, both of which play key osmoregulatory roles in land crabs. We (1) identified which genes were differentially expressed after exposure to desiccation stress in each species and across both tissue types, (2) investigated whether modules of orthologous genes with correlated expression across the transcriptome were more closely associated with a species’ terrestrial grade or with its relative phylogenetic distance to either of its confamilials, and (3) inferred the evolutionary histories of these genes (i.e., whether they were species-specific genes, single-copy conserved genes, or genes that have undergone gene expansion) and their putative functions.

In terms of overall tissue-specific patterns of gene expression, our DESeq2 differential expression analyses showed that *T. celeste* and *G. natalis* responded very strongly in their posterior gills, whereas *T. magna* responded to a lesser degree in its antennal gland. Since posterior gills in land crabs play more of an osmoregulatory role than a respiratory one (Péqueux 1995), this finding suggests that both *T. celeste* and *G. natalis* were under significant osmoregulatory stress 24h after the start of the desiccation stress trial. In fact, the greatest gene expression response for both *T. celeste* and *G. natalis*, both in terms of quantities of DEGs and magnitude of expression changes, occurred in their posterior gills during the Baseline – Acute time interval. In contrast, *T. magna* displayed significantly fewer DEGs than the other two species during the same time interval in its gills (Fig. 3C and Fig. 3D). But interestingly, for both *T. magna* and *G. natalis* at this time interval, nearly all differentially expressed genes that matched to functionally informative NCBI records were related to ATP generation and protein synthesis and were strongly upregulated (Table 2), implying that the acute response to desiccation for both species may utilize many energy intensive cellular processes even though the overall magnitude of their respective responses varied. This finding is consistent with another study that examined mitogenome evolution in panpulmonate gastropods (*i.e*., terrestrial snails and slugs), that revealed signatures of positive selection in genes related to oxidative phosphorylation (*cob* and *nad5*), suggesting that higher metabolic requirements on land may lead to an increased demand for cellular energy (Romero et al. 2016).

Interpreting these patterns rests on understanding whether *T. celeste* and *G. natalis* may have been in fact responding to different kinds of osmoregulatory stress. *T. celeste* relies heavily on freshwater to release its nitrogenous waste, and cannot release this waste unless its gills are submerged (Dela-Cruz and Morris 1997). Consequently, changes in posterior gill gene expression may truly represent a response to the acute desiccation stress experienced at the start of the time trial for this species. *G. natalis*, however, is highly terrestrial and inhabits dry burrows that do not contact the water table. It uses thick setal tufts on the coxae of its posterior walking legs to passively wick moisture from dew drops and puddles (Watson-Zink 2021). Since these tufts were constantly submerged during the 5d acclimation period, and four otherwise healthy *G. natalis* crabs unexpectedly perished during acclimation, the acclimation conditions themselves may actually have been stressful for this species, to the point that the crabs succumbed to overhydration and extreme hypo-osmotic stress. In this instance, the gene expression signal from its gills 24h after the start of the desiccation trial may still reflect its extreme stress response during the so-called acclimation period.

Since *T. celeste* relies on submersion in freshwater for urination (Weihrauch et al. 2004; Dela-Cruz and Morris 1997), we expected to observe more of a response in this species during the Extreme – Recovery time interval. But compared to its response during the Baseline – Acute time interval in its posterior gills, its response during the Extreme – Recovery time interval was relatively muted (3,503 vs 82 DEGs, respectively). Because *T. celeste* rapidly releases its nitrogenous wastes within 20 minutes to an hour after reimmersion (Dela-Cruz and Morris 1997), the possibility remains that the sampling 24h post re-immersion failed to capture the majority of this recovery response.

Several genes known to play critical roles in osmoregulation and excretion appeared to be differentially expressed in response to desiccation. But many of these genes (specifically the sodium-potassium ATPase, the sodium bicarbonate cotransporter, and the Rh-like protein) were downregulated at the start of the experiment and remained “off” throughout the length of the challenge for *T. celeste*, the least terrestrially adapted land crab in this experiment. This was also the case for an aquaporin gene in *G. natalis*. These findings were surprising, as other studies that have examined the genetic basis of terrestriality found that genes related to nitrogenous waste metabolism (in walking catfishes; Li et al. 2018) and ammonia excretion pathways (in mudskippers; You et al. 2014) played critical roles in allowing these species adapt to land. Aquaporins may also be especially important in enabling ancestrally marine animals to inhabit terrestrial environments. In vertebrates alone, there are 17 classes of aquaporins (Finn et al. 2014), and more generally across the Tree of Life, parallel aquaporin-coding gene duplications have been identified more often in animals that have transitioned from marine to non-marine environments (Martinez-Redondo et al. 2023). These findings may suggest that there may be other genes in the land crabs that play larger roles in helping these animals resist desiccation.

After grouping differentially expressed genes into modules of similar expression, differences between species best explained the prevailing signal. This may simply reflect the estimated 700,000 years of independent evolution between *T. celeste* and *T. magna* (Ng and Shih 2014), and presumably the even greater phylogenetic distance between the *Tuerkayana* crabs and their confamilial, *G. natalis*. The patterns in Modules 1 and 3 were consistent with shared ancestry (i.e., more similarities in gene expression responses between *T. celeste* and *T. magna*, which are sister species), implying that these particular modules may be under greater degrees of phylogenetic constraint. In other cases (for Modules 5 and 6), land crabs from similar terrestrial grades (i.e., *T. magna* and *G. natalis*, which are confamilial species) responded more similarly. This suggests that genes in these modules may either be related to genes that allow these crabs to display greater degrees of terrestriality (as may be the case for Module 6), or alternatively, related to genes that allow *T. celeste* to survive in its unique terrestrial niche, restricted to the shores of freshwater streams for urinary purposes, (as may be the case for Module 5). Interestingly, for Module 5, *T. celeste* showed a U-shaped response curve in both its antennal gland and its posterior gill, reflecting the overall degree of protein translation for genes in this module through time. Expression levels dropped significantly after the start of the desiccation trial for this species and began to increase again post-recovery, suggesting that desiccation stress causes *T. celeste* to shut down protein translation for genes in this module until its hydration status improves. In the posterior gill in particular, during the Acute – Extreme time interval, modular gene expression closely resembled that seen for *T. magna* and *G. natalis*, suggesting adaptive plasticity in response to hydration status in this species.

The single-copy conserved genes in the “terrestrial grade effect” Modules 5 and 6 primarily encode transcription factors for cellular protein complexes and chaperones. Species-level differences in gene expression for transcription factors and chaperones potentially reflect constitutive differences in protein translation for specific protein complexes between species. While an in-depth analysis of these key complexes and the involved biochemical pathways is outside the scope of this study, future work should determine whether these pathways allow land crabs to mitigate the osmoregulatory challenges associated with their terrestrial lifestyles.

Finally, while DEGs tended to be conserved genes, the DEGs with the greatest degree of differential expression for all three species were exclusively genes that were part of expanded gene families or novel genes that were not assigned to any orthogroup. These findings are consistent with those seen in terrestrial gastropods, which have colonized land at least 30 times across various ecological routes (Vermeij and Watson-Zink 2022). In gastropods, parallel expansions of ancient gene families critically enable mollusks to transition to land, suggesting that a pre-existing genomic “toolkit” may allow animals to adapt to novel habitats. However, lineage-specific gene gains simultaneously facilitated these transitions in gastropods (Aristide and Fernandez 2023), which also appears to be the case in the land crabs examined in this study. In *T. celeste*, 85% of its gill-derived transcriptomic response to acute desiccation stress was driven by novel, species-specific genes (Fig. 6). Furthermore, in *T. celeste*, novel genes were overwhelmingly upregulated in response to acute desiccation stress, whereas single-copy, conserved genes were significantly downregulated, which further supports the hypothesis that this species primarily uses novel genes to combat desiccation stress while on land.

Our conclusions regarding the evolutionary histories of the differentially expressed genes should be viewed against the background of using fragmented and incomplete transcriptome assemblies, leaving open the possibility that we may have overestimated the number of genes we recovered in our expanded gene families, or potentially missed genes with low expression altogether. Nevertheless, the results lay the groundwork for future studies using whole genome phylogenomic approaches to explore the role that gene repertoire evolution (i.e., gains, duplications, and losses) plays in the repeated convergent terrestrialization of the land crabs.

## Acknowledgements

We thank A. Whitehead and L. O’Connell for advice on experimental design, data analysis, and for constructive comments on earlier versions of the manuscript, the park rangers at the Christmas Island National Park for their assistance and fieldwork support, P.K.L. Ng for logistical support/international crab acquisition, S.A. Caplins and B. Cameron for advice on laboratory and bioinformatic methodologies, and all land crabs that were sacrificed to generate the data upon which this work is based.

