## supplemental materials for "Transcriptomic responses of gecarcinid land crabs to acute and prolonged desiccation stress"

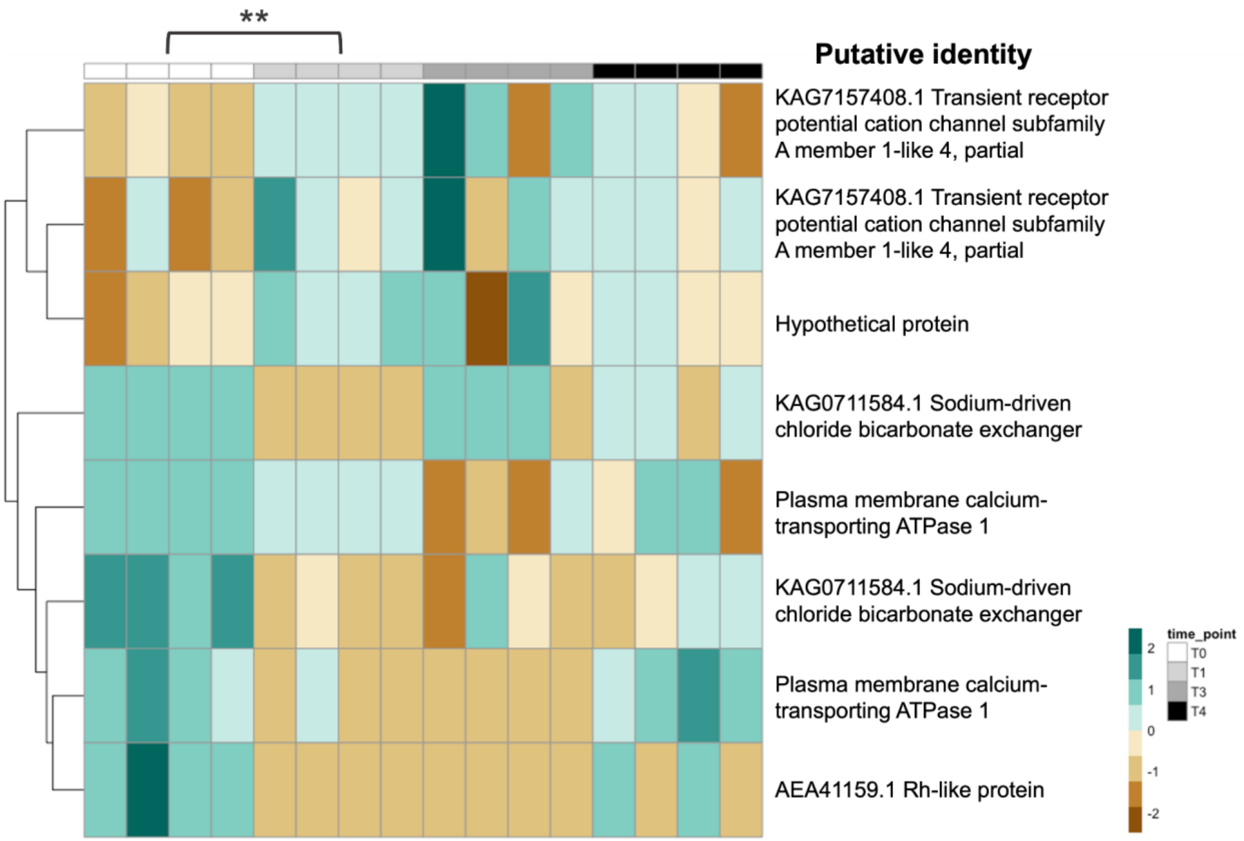


**Supplemental Materials, Figure 1:** Heatmap showing gene expression profiles for candidate genes displaying significant differential expression levels during experimental period in posterior gills of *T. celeste*. Asterisks reflect degree of statistical significance between log2FoldChange expression values between time points: $p \leq 0.05$ (*); $p \leq 0.01$ (**); $p \leq 0.001$ (***).

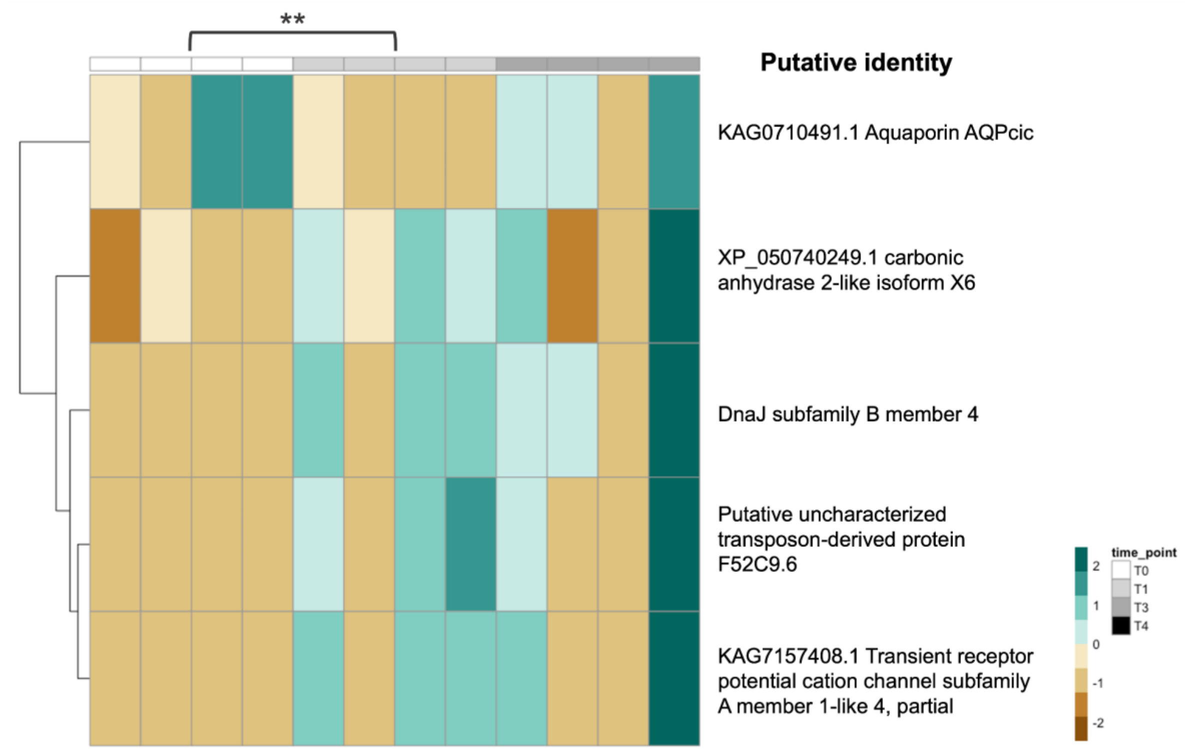


**Supplemental Materials, Figure 2:** Heatmap showing gene expression profiles for candidate genes displaying significant differential expression levels during experimental period in posterior gills of *G. natalis*. Asterisks reflect degree of statistical significance between log2FoldChange expression values between time points: $p \leq 0.05$ (*); $p \leq 0.01$ (**); $p \leq 0.001$ (***).

***T. celeste***

| | Antennal Gland | | | | | | Posterior Gill | | | | | |
|---|---|---|---|---|---|---|---|---|---|---|---|---|
| | | Expanded | Single-copy | | Unassigned | Assigned | | Expanded | Single-copy | | Unassigned | Assigned |
| Baseline - Acute | DEG | 19 | 0 | DEG | 149 | 19 | DEG | 291 | 235 | DEG | 2936 | 526 |
| | nonDEG | 112 | 3 | nonDEG | 17509 | 115 | nonDEG | 158 | 65 | nonDEG | 11280 | 223 |
| | odds ratio | | inf | odds ratio | | 0.111 | odds ratio | | 0.459 | odds ratio | | 0.151 |
| | adj. p-value | | 0.493 | adj. p-value | | **3.34E-08** | adj. p-value | | **3.28E-06** | adj. p-value | | **7.55E-101** |
| | | Expanded | Single-copy | | Unassigned | Assigned | | Expanded | Single-copy | | Unassigned | Assigned |
| Acute - Extreme | DEG | 5 | 1 | DEG | 62 | 6 | DEG | 6 | 3 | DEG | 34 | 9 |
| | nonDEG | 122 | 5 | nonDEG | 24604 | 127 | nonDEG | 50 | 3 | nonDEG | 8657 | 53 |
| | odds ratio | | 0.899 | odds ratio | | 0.05 | odds ratio | | 0.082 | odds ratio | | 0.024 |
| | adj. p-value | | 1 | adj. p-value | | **4.29E-11** | adj. p-value | | **1.14E-02** | adj. p-value | | **5.53E-10** |
| | | Expanded | Single-copy | | Unassigned | Assigned | | Expanded | Single-copy | | Unassigned | Assigned |
| Extreme - Recovery | DEG | 9 | 2 | DEG | 35 | 11 | DEG | 5 | 1 | DEG | 69 | 6 |
| | nonDEG | 185 | 45 | nonDEG | 25751 | 230 | nonDEG | 58 | 4 | nonDEG | 9689 | 62 |
| | odds ratio | | inf | odds ratio | | 0.076 | odds ratio | | inf | odds ratio | | 0.25 |
| | adj. p-value | | 0.288 | adj. p-value | | **8.37E-06** | adj. p-value | | 1 | adj. p-value | | 0.145 |

***T. magna***

| | Antennal Gland | | | | | | Posterior Gill | | | | | |
|---|---|---|---|---|---|---|---|---|---|---|---|---|
| | | Expanded | Single-copy | | Unassigned | Assigned | | Expanded | Single-copy | | Unassigned | Assigned |
| Baseline - Acute | DEG | 47 | 13 | DEG | 202 | 60 | DEG | 11 | 4 | DEG | 53 | 15 |
| | nonDEG | 237 | 115 | nonDEG | 20681 | 352 | nonDEG | 73 | 3 | nonDEG | 9885 | 76 |
| | odds ratio | | 1.05 | odds ratio | | 0.093 | odds ratio | | 0.187 | odds ratio | | 0.046 |
| | adj. p-value | | 1 | adj. p-value | | **2.50E-18** | adj. p-value | | 0.079 | adj. p-value | | **4.02E-12** |
| | | Expanded | Single-copy | | Unassigned | Assigned | | Expanded | Single-copy | | Unassigned | Assigned |
| Acute - Extreme | DEG | 23 | 3 | DEG | 84 | 26 | DEG | 10 | 1 | DEG | 28 | 11 |
| | nonDEG | 117 | 2 | nonDEG | 18126 | 119 | nonDEG | 152 | 25 | nonDEG | 13223 | 177 |
| | odds ratio | | 0.62 | odds ratio | | 0.053 | odds ratio | | 0.718 | odds ratio | | 0.038 |
| | adj. p-value | | 0.75 | adj. p-value | | **1.92E-15** | adj. p-value | | 0.677 | adj. p-value | | **4.23E-11** |
| | | Expanded | Single-copy | | Unassigned | Assigned | | Expanded | Single-copy | | Unassigned | Assigned |
| Extreme - Recovery | DEG | 11 | 2 | DEG | 38 | 13 | DEG | 23 | 3 | DEG | 63 | 26 |
| | nonDEG | 125 | 3 | nonDEG | 19077 | 128 | nonDEG | 77 | 5 | nonDEG | 10436 | 82 |
| | odds ratio | | inf | odds ratio | | 0.059 | odds ratio | | 0.2 | odds ratio | | 0.028 |
| | adj. p-value | | 0.493 | adj. p-value | | **5.42E-08** | adj. p-value | | 0.119 | adj. p-value | | **5.15E-23** |

***G. natalis***

| | Antennal Gland | | | | | | Posterior Gill | | | | | |
|---|---|---|---|---|---|---|---|---|---|---|---|---|
| | | Expanded | Single-copy | | Unassigned | Assigned | | Expanded | Single-copy | | Unassigned | Assigned |
| Baseline - Acute | DEG | 13 | 1 | DEG | 61 | 14 | DEG | 213 | 252 | DEG | 1261 | 465 |
| | nonDEG | 392 | 106 | nonDEG | 16622 | 498 | nonDEG | 27 | 2 | nonDEG | 4363 | 29 |
| | odds ratio | | 0.873 | odds ratio | | 0.109 | odds ratio | | 0.053 | odds ratio | | 0.053 |
| | adj. p-value | | 1 | adj. p-value | | **1.78E-05** | adj. p-value | | **1.80E-21** | adj. p-value | | **1.54E-155** |
| | | Expanded | Single-copy | | Unassigned | Assigned | | Expanded | Single-copy | | Unassigned | Assigned |
| Acute - Extreme | DEG | 18 | 7 | DEG | 95 | 25 | DEG | 0 | 0 | DEG | 0 | 0 |
| | nonDEG | 671 | 202 | nonDEG | 18511 | 873 | nonDEG | 59 | 8 | nonDEG | 5373 | 67 |
| | odds ratio | | 0.737 | odds ratio | | 0.104 | odds ratio | | inf | odds ratio | | 0 |
| | adj. p-value | | 0.628 | adj. p-value | | **1.31E-15** | adj. p-value | | 1 | adj. p-value | | 0.053 |

**Supplemental Materials, Table 1:** Fisher tests were performed on both tissue types for all three species at all three time intervals for (1) associations between differential expression status and whether genes were part of expanded gene families or were single-copy, and (2) associations between differential expression status and assignment status (*i.e.*, assigned to orthogroup or unique to that particular species). The p-values that were statistically significant are bolded.

| Tissue | Time interval | Target accession | Sequence identity | e-value | Length | log2FoldChange | Protein name | Description | Gene type | Copy number (TC) | Copy number (TM) | Copy number (GN) |
|---|---|---|---|---|---|---|---|---|---|---|---|---|
| AN | T0 - T1 | MPC39525.1 | 89.3 | 2.70E-116 | 225 | 10.9817307 | Protein kinase C | Cellular signal transduction | EX | 5 | 2 | 1 |
| AN | T1 - T3 | KAF7655963.1 | 38.9 | 2.40E-13 | 126 | 8.723233298 | hypothetical protein LDENG_00047990 | | U | | | |
| AN | T1 - T3 | MPC19370.1 | 86.7 | 0.00033 | 30 | -7.732305729 | Adrenodoxin, mitochondrial | Reduces mitochondrial cytochrome P450 for steroidogenesis | U | | | |
| AN | T1 - T3 | XP_042231732.1 | 86.7 | 4.80E-52 | 113 | -7.412235051 | chloride intracellular channel exl-1-like | Probable chloride channel | U | | | |
| AN | T3 - T4 | MPC46755.1 | 56.9 | 1.10E-08 | 51 | -8.813210662 | hypothetical protein E2C01_04048 | | EX | 4 | 1 | 0 |
| AN | T3 - T4 | QBA18592.1 | 80 | 0 | 826 | 8.288520041 | domeless | Epithelial morphogenesis during oogenesis; border cell migration | EX | 2 | 1 | 5 |
| GL | T0 - T1 | MPC28031.1 | 100 | 1.70E-20 | 50 | 9.776347529 | 60S ribosomal protein L39 | Protein synthesis | U | | | |
| GL | T1 - T3 | KAG0723700.1 | 94.8 | 4.60E-132 | 248 | 7.430899999 | Ubiquitin recognition factor in ER-associated degradation protein 1 | degrades ubiquitin fusion proteins | EX | 2 | 2 | 2 |
| GL | T1 - T3 | KAF7655963.1 | 38.9 | 2.40E-13 | 126 | 7.309283009 | hypothetical protein LDENG_00047990 | | U | | | |
| GL | T3 - T4 | KAG0730152.1 | 64.2 | 2.50E-16 | 81 | -8.838279507 | hypothetical protein GWK47_028855 | | U | | | |

**Supplemental Materials, Table 2**: Gene names and descriptions for top 10 DEGs in terms of log2FoldChange expression for *T. celeste* for both tissues (GL: Posterior gill 7; AN: Antennal gland) and all time intervals (T0 – T1: Baseline – Acute; T1 – T3: Acute – Extreme; T3 – T4: Extreme – Recovery). Only genes with BLAST hits are shown. Gene types (EX: expanded gene family; U: unassigned to any orthogroup), and copy numbers per species (TC: *T. celeste*; TM: *T. magna*; GN: *G. natalis*) are also shown.

| Tissue | Time interval | Target accession | Sequence identity | e-value | Length | log2FoldChange | Protein name | Description | Gene type | Copy number (TC) | Copy number (TM) | Copy number (GN) |
|---|---|---|---|---|---|---|---|---|---|---|---|---|
| AN | T0 - T1 | MPC48966.1 | 92.1 | 9.70E-61 | 126 | 9.77458949 | Multiple coagulation factor deficiency protein 2 | Secretion of coagulation factors | EX | 2 | 8 | 1 |
| AN | T0 - T1 | XP_034046538.1 | 70.8 | 1.30E-15 | 72 | -9.606411379 | tigger transposable element-derived protein 1-like | Part of pogo DNA-mediated transposon family | U | | | |
| AN | T0 - T1 | KAG0712991.1 | 67 | 3.50E-167 | 579 | 9.587521266 | Inactive serine/threonine-protein kinase TEX14 | Required both for the formation of intercellular bridges during meiosis and for kinetochore-microtubule attachment during mitosis | EX | 1 | 4 | 0 |
| AN | T0 - T1 | ROT64668.1 | 91.2 | 1.10E-247 | 453 | 9.090942593 | hypothetical protein C7M84_017386 | | EX | 2 | 5 | 1 |
| AN | T1 - T3 | KAG0713905.1 | 61 | 8.10E-23 | 118 | 10.36062669 | hypothetical protein GWK47_015179 | | U | | | |
| AN | T1 - T3 | MPC60099.1 | 94.3 | 2.20E-18 | 53 | 9.438728697 | hypothetical protein E2C01_054136 | | U | | | |
| AN | T1 - T3 | KAG0712991.1 | 67 | 3.50E-167 | 579 | 9.308795985 | Inactive serine/threonine-protein kinase TEX14 | Required both for the formation of intercellular bridges during meiosis and for kinetochore-microtubule attachment during mitosis | EX | 1 | 4 | 0 |
| AN | T1 - T3 | XP_042239947.1 | 96.3 | 6.00E-41 | 82 | 8.996840343 | transcription elongation factor 1 homolog | Maintains chromatin structure in actively transcribed regions | U | | | |
| AN | T3 - T4 | MPC60099.1 | 94.3 | 2.20E-18 | 53 | -9.20474339 | hypothetical protein E2C01_054136 | | U | | | |
| AN | T3 - T4 | KAG0703349.1 | 90.9 | 2.90E-188 | 363 | -8.929119267 | Fructose-bisphosphate aldolase | Plays key role in glycolysis | EX | 1 | 6 | 0 |
| AN | T3 - T4 | XP_042239782.1 | 86.5 | 1.20E-222 | 593 | 8.664685394 | poly(U)-binding-splicing factor PUF60-like isoform X3 | DNA- and RNA-binding protein, involved in several nuclear processes such as pre-mRNA splicing, apoptosis and transcription regulation | EX | 7 | 6 | 8 |
| AN | T3 - T4 | KAG0723842.1 | 90.6 | 3.40E-206 | 385 | 8.188012272 | Beta-ureidopropionase | Catalyzes a late step in pyrimidine degradation | U | | | |
| GL | T0 - T1 | KAG0722361.1 | 76.5 | 2.20E-44 | 119 | -8.375777505 | RNA-directed DNA polymerase from mobile element jockey | Reverse transcriptase | U | | | |
| GL | T0 - T1 | CAR82606.1 | 96.8 | 5.80E-122 | 218 | 8.351752235 | mitochondrial manganese superoxide dismutase | Mitigates reactive oxygen species in the mitochondrial matrix to prevent mitochondrial dysfunction | EX | 3 | 8 | 1 |
| GL | T0 - T1 | KAG6654567.1 | 67.6 | 0.012 | 37 | 8.140118316 | hypothetical protein CIPAW_05G154200 | | U | | | |
| GL | T0 - T1 | KAG0711760.1 | 39.8 | 1.10E-20 | 226 | 8.08762212 | hypothetical protein GWK47_019943 | | EX | 9 | 9 | 1 |
| GL | T1 - T3 | XP_042218110.1 | 97.4 | 2.20E-111 | 228 | -9.689122289 | POU domain, class 6, transcription factor 2-like | Probable transcription factor likely to be involved in early steps in the differentiation of amacrine and ganglion cells | U | | | |
| GL | T1 - T3 | ROT85526.1 | 86.5 | 0.00018 | 37 | 8.890028557 | putative ribosome-binding protein 1-like isoform X11 | ribosome receptor and mediates interaction between the ribosome and the endoplasmic reticulum membrane | EX | 2 | 14 | 3 |
| GL | T1 - T3 | MPC17943.1 | 94.8 | 0 | 659 | 8.516994934 | Stress-70 protein, mitochondrial | Chaperone protein which plays an important role in mitochondrial iron-sulfur cluster (ISC) biogenesis | EX | 1 | 3 | 1 |
| GL | T1 - T3 | KAG0729092.1 | 95.1 | 1.90E-171 | 385 | -8.323543751 | Hippocampus abundant transcript 1 protein | Involved in transmembrane transport | EX | 2 | 2 | 2 |
| GL | T1 - T3 | MPC29685.1 | 91 | 1.10E-179 | 368 | -8.182291567 | hypothetical protein E2C01_022931 | | EX | 2 | 5 | 1 |
| GL | T3 - T4 | XP_027214065.1 | 87.4 | 2.10E-83 | 175 | 9.061012115 | remodeling and spacing factor 1-like | Regulatory subunit of the ATP-dependent RSF-1 and RSF-5 ISWI chromatin-remodeling complexes | EX | 0 | 2 | 1 |
| GL | T3 - T4 | KAG0695240.1 | 93.2 | 1.90E-173 | 340 | -8.552995795 | DNA repair protein RAD51 1 | Required both for recombination and for the repair of DNA damage | EX | 1 | 3 | 1 |
| GL | T3 - T4 | XP_042242848.1 | 97.4 | 9.00E-12 | 190 | 8.222477714 | ankyrin-2-like isoform X13 | Enables ATPase binding activity; potassium channel regulator activity; and transmembrane transporter binding activity | EX | 22 | 2 | 5 |
| GL | T3 - T4 | ATD53652.1 | 81.8 | 1.70E-241 | 499 | 8.055246545 | scavenger receptor class B1 | Receptor for different ligands such as phospholipids, cholesterol ester, lipoproteins, phosphatidylserine and apoptotic cells | EX | 7 | 5 | 5 |

**Supplemental Materials, Table 3**: Gene names and descriptions for top 10 DEGs in terms of log2FoldChange expression for *T. magna* for both tissues (GL: Posterior gill; AN: Antennal gland) and all time intervals (T0 – T1: Baseline – Acute; T1 – T3: Acute – Extreme; T3 – T4: Extreme – Recovery). Only genes with BLAST hits are shown. Gene types (EX: expanded gene family; U: unassigned to any orthogroup), and copy numbers per species (TC: *T. celeste*; TM: *T. magna*; GN: *G. natalis*) are also shown.

| Tissue | Time interval | Target accession | Sequence identity | e-value | Length | log2FoldChange | Protein name | Description | Gene type | Copy number (TC) | Copy number (TM) | Copy number (GN) |
|---|---|---|---|---|---|---|---|---|---|---|---|---|
| AN | T0 - T1 | KAG0722448.1 | 90.7 | 2.90E-220 | 408 | 23.40201232 | Tubulin polyglutamylase TTLL4 | Involved in KLF4 glutamylation leading to somatic cell reprogramming, pluripotency maintenance and embryogenesis | EX | 2 | 2 | 4 |
| AN | T0 - T1 | MPC13266.1 | 66 | 4.80E-92 | 1059 | 11.22971112 | hypothetical protein E2C01_005993 | | EX | 3 | 2 | 3 |
| AN | T0 - T1 | KAG0725138.1 | 72.4 | 6.60E-83 | 225 | 8.150209532 | Transcription factor A, mitochondrial | Transcription termination factor | U | | | |
| AN | T1 - T3 | XP_042204822.1 | 94.5 | 5.30E-67 | 201 | 24.86388684 | zinc finger matrin-type protein 2-like | Involved in pre-mRNA splicing as a component of the spliceosome. | EX | 1 | 4 | 4 |
| AN | T1 - T3 | KAG0716941.1 | 73.1 | 1.40E-127 | 327 | 23.48642385 | hypothetical protein GWK47_008452 | | EX | 3 | 1 | 5 |
| AN | T1 - T3 | KAG7172599.1 | 45 | 0.00096 | 60 | 23.07693708 | hypothetical protein Hamer_G006808 | | U | | | |
| AN | T1 - T3 | WP_208094023.1 | 72.9 | 5.40E-84 | 310 | 13.81598152 | hypothetical protein | | EX | 6 | 5 | 2 |
| GL | T0 - T1 | YP_009107263.1 | 77.9 | 5.50E-27 | 95 | 12.86102952 | NADH dehydrogenase subunit 5 | Part of mitochondrial respiratory chain complex I | U | | | |
| GL | T0 - T1 | YP_009535655.1 | 84.1 | 8.80E-86 | 208 | 11.6190675 | cytochrome c oxidase subunit 1 | Component of the cytochrome c oxidase, the last enzyme in the mitochondrial electron transport chain which drives oxidative phosphorylation. | U | | | |
| GL | T0 - T1 | KPV71458.1 | 90 | 1.60E-12 | 40 | 11.6177469 | hypothetical protein RHOBADRAFT_19411 | | U | | | |
| GL | T0 - T1 | OFQ97201.1 | 94 | 2.20E-15 | 50 | 11.5101051 | hypothetical protein HMPREF2909_08580 | | U | | | |
| GL | T0 - T1 | QRK27379.1 | 84.9 | 2.10E-21 | 73 | 10.75255852 | NADH dehydrogenase subunit 4 | Part of mitochondrial respiratory chain complex I | U | | | |
| GL | T0 - T1 | ORD93174.1 | 96.5 | 1.20E-41 | 85 | 10.29680205 | hypothetical protein ECANGB1_812 | | U | | | |
| GL | T0 - T1 | YP_009535658.1 | 79.1 | 5.80E-53 | 158 | 10.12406695 | ATP synthase F0 subunit 6 | Key component of the proton channel; it plays a direct role in the translocation of protons across the membrane | U | | | |
| GL | T0 - T1 | YP_009535661.2 | 80.2 | 1.70E-28 | 91 | 9.910830137 | NADH dehydrogenase subunit 5 | Part of mitochondrial respiratory chain complex I | U | | | |

**Supplemental Materials, Table 4**: Gene names and descriptions for top 10 DEGs in terms of log2FoldChange expression for *G. natalis* for both tissues (GL: Posterior gill; AN: Antennal gland) and all time intervals (T0 – T1: Baseline – Acute; T1 – T3: Acute – Extreme). Only genes with BLAST hits are shown. Gene types (EX: expanded gene family; U: unassigned to any orthogroup), and copy numbers per species (TC: *T. celeste*; TM: *T. magna*; GN: *G. natalis*) are also shown.

**Supplemental Materials, Table 5:** Gene names and descriptions for top 10 genes in terms of Module Membership for all modules with significant correlations to species. Only genes with BLAST hits are shown.

| Module | Target accession | Protein name | Description | Length | Sequence identity | e-value | \|Module Membership\| |
|---|---|---|---|---|---|---|---|
| ME1 | XP_042216344.1 | fumarylacetoacetase-like | phenylalanine and tyrosine degradation | 416 | 80 | 3.90E-206 | 0.977858078 |
| | KAG0725090.1 | Elongin-C | subunit of the transcription factor B (SIII) complex | 117 | 99.1 | 3.20E-60 | 0.971496157 |
| | MPC37806.1 | hypothetical protein | | 148 | 81.1 | 1.00E-56 | 0.971187492 |
| | ROT79510.1 | RMD5-like protein A | negative regulation of gluconeogenesis; ubiquitin protein ligase activity | 263 | 93.9 | 1.20E-139 | 0.956820355 |
| | KAG0724140.1 | Endoribonuclease LACTB2 | RNA phosphodiester bond hydrolysis | 302 | 68.2 | 2.80E-114 | 0.954221121 |
| | KAG0704322.1 | Ras-like GTP-binding protein Rho1 | actin cytoskeletal organization | 194 | 99 | 1.90E-104 | 0.952294347 |
| | XP_042219201.1 | short/branched chain specific acyl-CoA dehydrogenase, mitochondrial-like isoform X1 | fatty acid metabolic process; isoleucine catabolic process | 418 | 83 | 3.00E-193 | 0.950659183 |
| | AIS24843.1 | T-complex protein 1 subunit alpha | TRiC complex mediates the folding of WRAP53/TCAB1, thereby regulating telomere maintenance | 555 | 95 | 1.40E-292 | 0.949287268 |
| | MPC23252.1 | RPII140-upstream gene protein-like | RNA polymerase | 278 | 64.7 | 1.30E-94 | 0.947865926 |
| | XP_037797882.1 | phytanoyl-CoA dioxygenase, peroxisomal-like isoform X2 | fatty acid metabolism | 296 | 79.7 | 8.40E-145 | 0.945516155 |
| ME3 | KAG0724012.1 | COP9 signalosome complex subunit 6 | protein deneddylation | 293 | 96.6 | 1.70E-156 | 0.95228519 |
| | ALP46200.1 | nascent polypeptide-associated complex alpha | negative regulation of protein localization to endoplasmic reticulum | 209 | 100 | 1.00E-81 | 0.94535869 |
| | ROT63357.1 | grpE protein homolog 1, mitochondrial-like | controls the nucleotide-dependent binding of mitochondrial HSP70 to substrate proteins | 143 | 81.1 | 1.60E-58 | 0.93943885 |
| | XP_042231848.1 | cleavage and polyadenylation specificity factor subunit 5-like | activator of the pre-mRNA 3'-end cleavage and polyadenylation processing required for the maturation of pre-mRNA into functional mRNAs | 228 | 99.1 | 2.00E-129 | 0.93924806 |
| | ROT69872.1 | hypothetical protein | | 284 | 87.3 | 1.20E-139 | 0.93869091 |
| | KAG0717700.1 | U4/U6 small nuclear ribonucleoprotein Prp31 | Involved in pre-mRNA splicing as component of the spliceosome | 494 | 91.1 | 1.60E-243 | 0.93431374 |
| | KAG0723646.1 | Hydroxysteroid dehydrogenase-like protein 2 | oxidoreductase activity | 420 | 85.7 | 2.00E-204 | 0.92884357 |
| | XP_042210450.1 | trigger factor-like | Chaperone activity | 71 | 40.8 | 0.00028 | 0.9279747 |
| | XP_042215298.1 | 60S ribosomal protein L31-like | cytoplasmic translation | 125 | 94.4 | 1.70E-58 | 0.92657039 |
| | KAG0710068.1 | Short-chain specific acyl-CoA dehydrogenase, mitochondrial | aerobic process breaking down fatty acids into acetyl-CoA and allowing the production of energy from fats | 434 | 86.2 | 9.80E-206 | 0.92628632 |

| | | | | | | | |
|---|---|---|---|---|---|---|---|
| **ME4** | EGW03135.1 | Elongation factor 2 | essential factor for protein synthesis | 395 | 100 | 1.60E-232 | 0.979095787 |
| | XP_042236946.1 | L-2-hydroxyglutarate dehydrogenase, mitochondrial-like isoform X3 | oxidoreductase activity | 402 | 82.8 | 3.50E-202 | 0.967639558 |
| | XP_042226004.1 | transitional endoplasmic reticulum ATPase | Autophagy, DNA damage, DNA repair, Transport, Ubl conjugation pathway | 794 | 98.4 | 0 | 0.966802196 |
| | MPC32893.1 | Protein ROP | Transcription regulation | 356 | 94.7 | 5.00E-192 | 0.965047157 |
| | MPC14080.1 | Histone deacetylase complex subunit SAP18 | mRNA processing, mRNA splicing, Transcription, Transcription regulation | 150 | 97.3 | 3.40E-79 | 0.955379515 |
| | QDE54938.1 | eukaryotic translation initiation factor 3 subunit B, partial | Protein biosynthesis, Initiation factor, RNA-binding | 562 | 93.1 | 0 | 0.954118167 |
| | KAG0710311.1 | AP-2 complex subunit mu | Endocytosis, Protein transport, Transport, adaptor protein complex | 293 | 100 | 1.90E-163 | 0.952188669 |
| | KAG0713716.1 | CDGSH iron-sulfur domain-containing protein 1 | regulating maximal capacity for electron transport and oxidative phosphorylation | 110 | 84.5 | 6.30E-50 | 0.948170144 |
| | ACY66390.1 | FK506-binding protein 1A | immunoregulation and cellular processes involving protein folding and trafficking | 110 | 87.3 | 7.40E-49 | 0.947954257 |
| | MPC68061.1 | hypothetical protein | | 216 | 88 | 3.90E-98 | 0.944352237 |
| **ME5** | KAG0714100.1 | Tyrosyl-DNA phosphodiesterase 2 | DNA repair | 363 | 72.2 | 6.10E-145 | 0.940638747 |
| | KAG0725244.1 | T-complex protein 1 subunit delta | The TRiC complex mediates the folding of WRAP53/TCAB1, thereby regulating telomere maintenance | 534 | 95.1 | 1.50E-279 | 0.937633129 |
| | QHD64853.1 | heat shock protein 60 | heat inducible and act as a molecular chaperones, assisting in protein folding | 556 | 97.3 | 6.90E-294 | 0.932209271 |
| | XP_042230358.1 | reticulon-4-interacting protein 1 homolog, mitochondrial-like isoform X1 | Plays a role in the regulation of retinal ganglion cell (RGC) maturation and neurite outgrowth | 370 | 68.6 | 9.50E-145 | 0.918332399 |
| | KAG0726339.1 | Methylthioribose-1-phosphate isomerase | L-methionine biosynthesis via salvage pathway | 348 | 81.9 | 1.80E-158 | 0.91789729 |
| | ANN46488.1 | calnexin | Chaperone activity | 442 | 95.9 | 4.30E-168 | 0.914040218 |
| | MPC18378.1 | Centromere protein X | Cell cycle, Cell division, DNA damage, DNA repair, Mitosis | 65 | 76.9 | 7.10E-18 | 0.911495751 |
| | KAG0729193.1 | Growth arrest and DNA damage-inducible proteins-interacting protein 1 | Acts as a negative regulator of G1 to S cell cycle phase progression by inhibiting cyclin-dependent kinases | 264 | 62.9 | 7.90E-63 | 0.90744937 |
| | MPC17875.1 | Oligosaccharyltransferase complex subunit ostc-B | involved in protein glycosylation and protein modification | 150 | 94 | 1.10E-73 | 0.905814954 |
| | KAG0726480.1 | Mitochondrial import inner membrane translocase subunit TIM14 | Chaperone activity | 84 | 91.7 | 1.80E-34 | 0.903171201 |

| | | | | | | | |
|---|---|---|---|---|---|---|---|
| ME6 | XP_042209693.1 | 40S ribosomal protein SA-like | Cell adhesion, cytoplasmic translation, ribosomal small subunit assembly | 212 | 96.2 | 1.70E-112 | 0.969824031 |
| | ACG60901.1 | eukaryotic translation initiation factor 6 | stimulatory translation initiation factor downstream insulin/growth factors | 245 | 86.5 | 4.50E-116 | 0.961991376 |
| | XP_042222113.1 | 60S ribosomal protein L24-like | encodes a ribosomal protein that is a component of the 60S subunit | 105 | 94.3 | 7.10E-49 | 0.954621492 |
| | XP_037784230.1 | eukaryotic translation initiation factor 3 subunit E-like | Component of the eukaryotic translation initiation factor 3 (eIF-3) complex | 424 | 95.3 | 4.00E-232 | 0.95200711 |
| | XP_042233103.1 | transcription elongation factor S-II-like isoform X1 | part of complex required to increase the catalytic rate of RNA polymerase II transcription | 301 | 83.4 | 1.30E-110 | 0.943234722 |
| | KAG7162228.1 | Mediator of RNA polymerase II transcription subunit 21-like | a coactivator involved in the regulated transcription of nearly all RNA polymerase II-dependent genes. | 147 | 98 | 6.60E-70 | 0.939832243 |
| | MPC31537.1 | hypothetical protein | | 95 | 98.9 | 2.90E-40 | 0.931888824 |
| | MPC56548.1 | hypothetical protein | | 381 | 57.7 | 7.80E-85 | 0.929501031 |
| | ADE60733.1 | myosin essential light chain | regulating the actin-myosin interaction of smooth muscle | 147 | 98 | 1.90E-73 | 0.925459286 |
| | KAG0714221.1 | Mitochondrial-processing peptidase subunit alpha | Substrate recognition and binding subunit of the essential mitochondrial processing protease | 324 | 90.1 | 6.60E-171 | 0.916987419 |
